# CHIP, VCP, and Nucleolar Gatekeepers Remodel the Nucleolus into a Stress-Responsive Proteostasis Hub

**DOI:** 10.1101/2025.10.04.680426

**Authors:** Małgorzata Piechota, Lilla Biriczová, Konrad Kowalski, Fernando Aprile García, Marta Leśniczak-Staszak, Natalia A. Szulc, Witold Szaflarski, Ritwick Sawarkar, Wojciech Pokrzywa

## Abstract

The nucleolus, classically dedicated to ribosome biogenesis, also acts as a stress-responsive proteostasis hub. During heat shock, misfolded proteins accumulate in its granular component (GC), but whether this is passive or regulated has been unclear. We show that the ubiquitin ligase CHIP remodels the nucleolus into a reversible protein quality control (PQC) compartment by diverting HSP70 from refolding toward sequestration and transiently suppressing rRNA synthesis. This process requires the segregase VCP, which mediates ubiquitinated substrate flux and couples nucleolar PQC with nuclear and ER stress responses. Functional genomics identify NOL6 and WDR55 as intrinsic gatekeepers with opposing effects on sequestration. We further define three structural states - free-flow, peripheral, and sealed - that encode PQC capacity and recovery potential. Thus, nucleolar proteostasis is revealed as an actively regulated continuum, linking chaperones, ubiquitination, and ribosome biogenesis with global stress adaptation.

**Highlights:**

- CHIP remodels the nucleolus into a reversible protein quality control (PQC) compartment during heat stress
- VCP drives flux of ubiquitinated substrates and prevents premature resumption of rRNA synthesis
- NOL6 promotes, while WDR55 restricts, nucleolar storage of misfolded proteins
- Distinct nucleolar morphologies (free-flow, peripheral, sealed) encode PQC states
- Ubiquitination and deubiquitination act as switchable signals, not passive damage marks
- Nucleolar PQC is integrated with ISR, ER stress, and stress granule pathways

## INTRODUCTION

The nucleolus is a multifunctional membraneless organelle canonically responsible for ribosome biogenesis, coordinating rRNA transcription, processing, and subunit assembly within its tripartite architecture - the fibrillar centre (FC), dense fibrillar component (DFC), and granular component (GC)^1,2^. Beyond this biosynthetic role, the nucleolus has emerged as a dynamic stress-responsive compartment that transiently engages in protein quality control (PQC)^3^.

Under proteotoxic conditions such as heat shock, misfolded proteins accompanied by HSP70 accumulate within the GC, converting the nucleolus to a reversible storage that buffers cytoplasmic toxicity and preserves chaperone availability^4–6^. Proteomic studies have identified stress-induced nucleolar visitors, including unfolded substrates, chaperones, and ubiquitin modifiers, positioning the nucleolus as a proteostasis reserve. Recent reports describe amyloid-like assemblies and nucleolar caps that reorganise nucleolar architecture and may separate ribosome assembly sites^7,8^. Post-translational regulators such as SENP3, USP36, and K29-linked ubiquitin (Ub) chains further connect PQC to nucleolar function^9–11^. Morphological reorganisations, such as nucleolar stress bodies in *C. elegans*^12^ and ring-shaped nucleoli in mammalian cells, suggest that structural transitions correlate with stress states. However, the molecular logic linking nucleolar architecture, protein sequestration, and rRNA output remains poorly understood. Moreover, how nucleolar PQC integrates with wider stress networks including the integrated stress response (ISR), endoplasmic reticulum (ER) stress pathways, and cytoplasmic stress granules, remains an open question.

The E3 ubiquitin ligase CHIP (STUB1) is a conserved PQC factor that couples the chaperone network to the ubiquitin-proteasome system (UPS). Through its tetratricopeptide repeat (TPR) and U-box domains, CHIP binds HSP70 and HSP90 to ubiquitinate unfolded or chaperone-bound clients^13–15^. This interaction is reciprocal: CHIP can attenuate HSP70-mediated refolding, biasing outcomes toward ubiquitination and clearance^16,17^, while HSP70 can stabilise CHIP in an auto-inhibited state^18,19^. CHIP thus acts as a tunable regulator of its clients’ fate, but whether it also governs the storage and recovery of misfolded proteins within the nucleolus has not been tested. The AAA^+^ ATPase VCP/p97, another central PQC regulator, functions as a Ub-selective segregase that extracts ubiquitinated proteins from complexes, condensates, or membrane compartments to facilitate their recycling or proteasomal degradation^20–22^. Increasing evidence indicates that VCP functionally intersects with chaperone systems, particularly the HSP70-CHIP axis. In protein quality control, HSP70 attempts client refolding, CHIP biases outcomes toward ubiquitination, and VCP subsequently extracts the ubiquitinated clients for downstream processing^23,24^. More broadly, chaperones and segregases can act either sequentially or in coordination to dissolve protein assemblies, as illustrated by VCP-mediated extraction of ubiquitinated G3BP1 to promote stress granule disassembly^25^. Whether such interplay also extends to the nucleolus remains unknown: does VCP actively extract ubiquitinated substrates during stress recovery, or does it cooperate with CHIP and HSP70 to balance sequestration and clearance? Furthermore, could these classical PQC factors intersect with intrinsic ribosome biogenesis proteins to jointly shape nucleolar dynamics under stress?

Here, we address these gaps by asking whether classical PQC machinery and intrinsic ribosome biogenesis factors cooperate to actively remodel the nucleolus during heat stress. We show that CHIP attenuates HSP70 activity, promotes ubiquitin-dependent sequestration of misfolded proteins, and represses rRNA synthesis; that VCP supports the flux of ubiquitinated proteins and timely recovery; and that nucleolar factors such as NOL6 and WDR55 tune scaffold properties to balance storage and biosynthesis. Together, our findings establish the nucleolus as a protein-directed PQC hub that integrates folding, ubiquitination, sequestration, and ribosome biogenesis into a coordinated stress response.

## RESULTS

### CHIP and HSP70 Cooperate to Modulate Nucleolar Engagement and Clearance during Heat-Induced Proteotoxic Stress

To test whether CHIP engages nucleolar PQC under proteotoxic stress, we generated Flp-In compatible HeLa and HEK-293 lines stably expressing EGFP-CHIP (HeLa CHIP, HEK CHIP; Fig. S1A).

To verify that this behaviour is not restricted to engineered systems, we additionally examined transient EGFP-CHIP expression in MCF7 breast cancer cells. Upon heat shock (42 °C, 90-120 min) followed by 2 h recovery, EGFP-CHIP accumulated in nucleoli and was released during recovery across all cellular backgrounds (Fig. 1A-B, S1B-C). This redistribution was specific to heat; arsenite (oxidative stress), sorbitol (osmotic stress), thapsigargin (ER stress) or puromycin (translational stress) did not drive CHIP to nucleoli (Fig. S1C). Consistent with prior studies, HSP70 was also enriched in nucleoli after heat (Fig. 1C)^4,26–30^. Super-resolution Airyscan imaging with NPM1 (GC) and fibrillarin/FBL (DFC) markers showed CHIP predominantly in the GC, with weak overlap with DFC (Fig. 1D-E), in line with GC-linked storage of misfolded clients^2,4,6,30^. Pretreatment of cells with VER-155008 (VER), an ATP-competitive inhibitor of the HSP70/Hsc70 ATPase domain, before heat shock did not affect CHIP retention to GC compartment (Fig. 1E, S1D).

**Figure 1.**
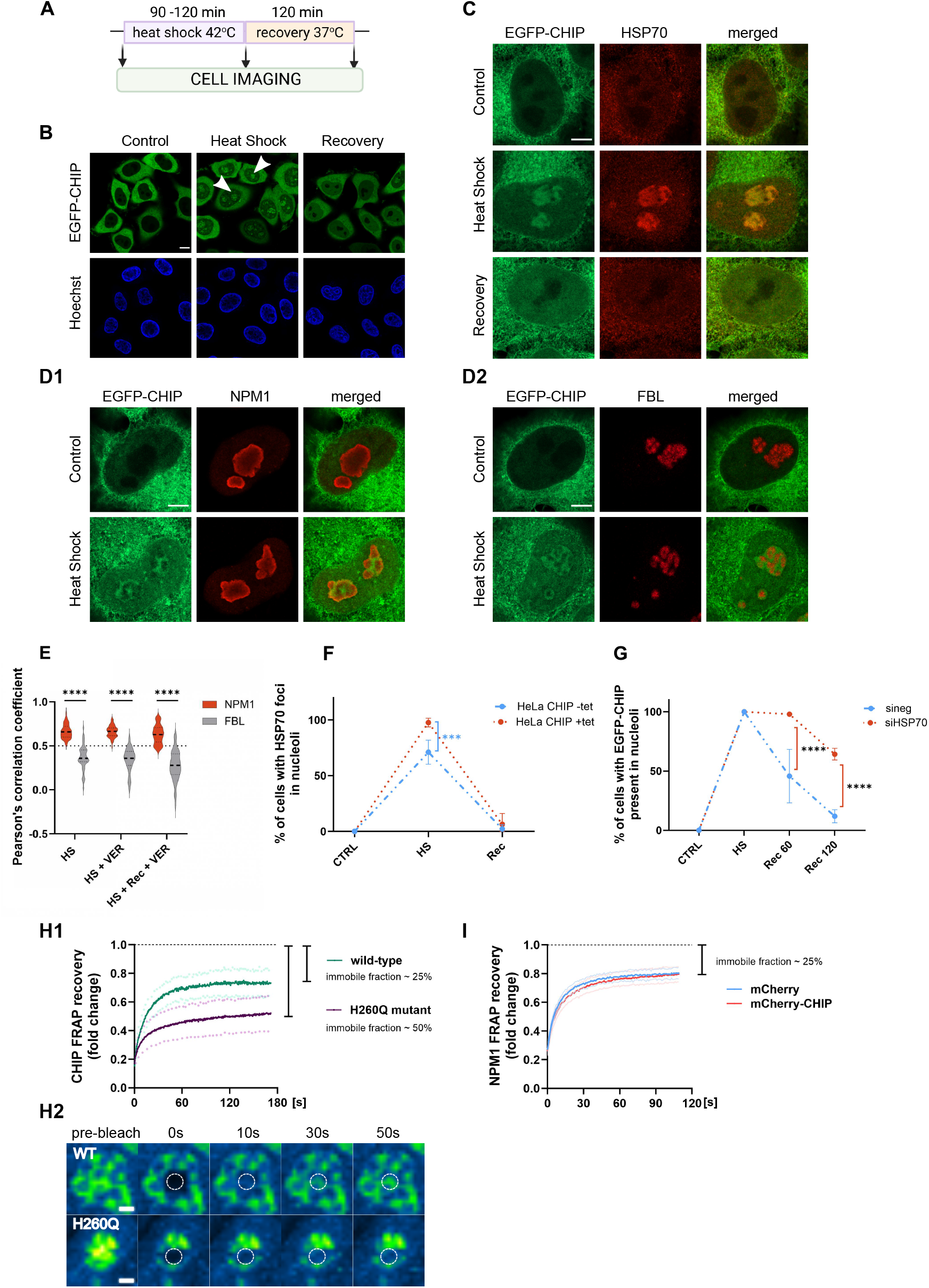
CHIP and HSP70 Cooperate to Control Nucleolar Engagement and Clearance during Heat Stress. (A) Schematic of the heat shock paradigm (42 °C, 90-120 min; recovery at 37 °C, 2 h unless indicated otherwise). (B) Representative confocal images of EGFP-CHIP in HeLa Flp-In cells under basal, heat shock, and recovery conditions. Arrowheads point at nucleoli. Scale bar, 10 µm. (C) Representative Airyscan images of HSP70 in HeLa CHIP cells after heat shock. Scale bar, 5 µm. (D) CHIP localisation within the nucleolar GC. (D1) EGFP-CHIP with NPM1 (GC marker). (D2) EGFP-CHIP with FBL (DFC marker). Scale bars, 5 µm. (E) Quantification of CHIP colocalisation with NPM1 or FBL. Violin plots of Pearson’s coefficient (n = 31–70 nucleoli). Median and quartiles shown. Unpaired two-tailed t-test. HS – heat shock, VER – VER-155008, Rec – recovery. (F) Percentage of cells with nucleolar HSP70 after heat shock and recovery with or without EGFP-CHIP induced by tetracycline (tet). Mean ± SD from n = 3 experiments. Two-way ANOVA with Sidak’s test. HS – heat shock, Rec – recovery. (G) CHIP nucleolar retention in cells transfected with HSP70 siRNA, (sineg - control siRNA). Mean ± SD from n = 3 experiments. Two-way ANOVA with Tukey’s test. HS – heat shock, Rec – recovery, the numbers next to it indicate the duration in minutes. (H) FRAP of EGFP-CHIP WT and H260Q mutant in nucleoli. (H1) Recovery curves (n = 9-12 nucleoli). Mean ± SD. (H2) Representative pseudo-coloured time-lapse images. Scale bars, 2 µm. (I) FRAP of EGFP-NPM1 with or without mCherry-CHIP. Recovery curves (n = 9-11 nucleoli). Mean ± SD. See also Figure S1

To probe functional crosstalk, we modulated HSP70 abundance and chaperone activity. Inducing CHIP (by tetracycline) increased nucleolar HSP70 across single cells (Fig. 1F), suggesting CHIP potentiates HSP70 engagement. Conversely, HSP70 depletion (siRNA; Fig. S1E) reduced CHIP entry (~40% decrease during heat shock) and strongly delayed its removal: ~60% of HSP70-depleted cells retained nucleolar CHIP at 2 h recovery versus ~10% in controls (Fig. 1G, S1F). We next used VER to acutely block chaperone cycling. VER partially phenocopied HSP70 depletion: CHIP entry was intact, but clearance was delayed (Fig. S1G), indicating that HSP70 ATP-dependent turnover is dispensable for CHIP recruitment but required for its disengagement.

To determine whether CHIP is dynamically engaged rather than passively retained, we performed FRAP (fluorescence recovery after photobleaching), in which a nucleolar ROI (region of interest) is bleached and signal recovery reports molecular exchange between the nucleolus and surrounding pools (mobile fraction and half-time). EGFP-CHIP recovered rapidly (Fig. 1H1-H2), whereas CHIP^H260Q^, a catalytically inactive mutant unable to perform ubiquitination^31^, showed markedly reduced recovery, consistent with substrate trapping or conformational immobilization. Importantly, CHIP overexpression did not alter NPM1 mobility in EGFP-NPM1 cells co-expressing mCherry-CHIP (as measured by FRAP), arguing that CHIP integration does not globally change nucleolar material properties (Fig. 1I). Together, these results indicate that CHIP engages with nucleoli dynamically, with its mobility shaped by catalytic activity and clearance strongly dependent on HSP70, consistent with an active disassembly mechanism during recovery.

### CHIP Expands Nucleolar Storage Capacity for Misfolded Proteins by Promoting HSP70-Mediated Sequestration

Having established that CHIP dynamically localises to nucleoli and cooperates with HSP70 (Fig. 1), we next examined whether it actively promotes the sequestration of misfolded proteins and influences recovery kinetics. Because nucleolar reorganisation often correlates with changes in FBL morphology, we first inspected FBL under stress. Heat shock reduced FBL levels, which were only partially restored after 2 h recovery, and inhibition of HSP70 ATPase activity with VER further impaired this re-accumulation (Fig. 2A). Co-labelling with ubiquitin (Ub) revealed a strong correlation between storage state and FBL distribution: nucleoli lacking Ub conjugates retained a compact, bright FBL signal, whereas Ub-positive nucleoli displayed reduced and dispersed FBL (Fig. 2B1, S2A). Notably, CHIP overexpression increased the fraction of nucleoli with dispersed FBL during heat shock and sustained this dispersed phenotype during recovery, preventing full restoration of compact morphology (Fig. 2B2). These findings suggest that CHIP enhances nucleolar storage capacity, but at the cost of re-establishing canonical nucleolar architecture.

**Figure 2.**
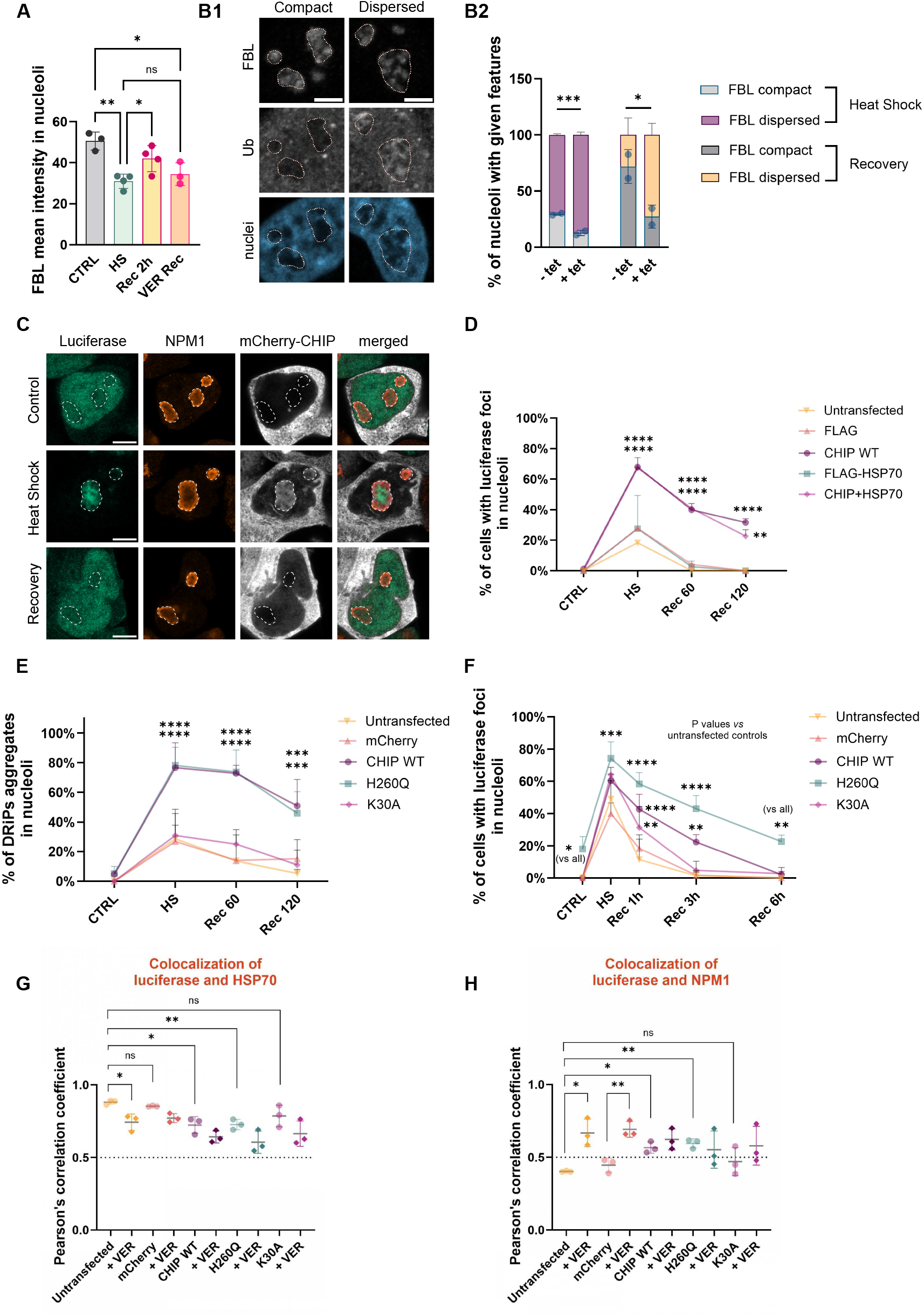
CHIP Expands Nucleolar Storage of Misfolded Proteins Through HSP70-mediated Sequestration. (A) Quantification of nucleolar FBL levels after heat stress and VER treatment. Mean ± SD from n = 3-4 experiments. One-way ANOVA with Tukey’s test. (B) CHIP overexpression increases the fraction of nucleoli with dispersed FBL after stress. (B1) Representative confocal images showing FBL and ubiquitin (Ub) staining in compact versus dispersed phenotypes. Scale bar, 5 µm. (B2) Quantification of nucleolar phenotypes after heat shock and recovery ± EGFP-CHIP induced by tetracycline (tet). Mean ± SD from n = 2 experiments. Two-way ANOVA with Fisher’s LSD test. (C) Confocal images of HEK293T cells expressing luciferase and mCherry-CHIP, immunostained for NPM1. Scale bar, 5 µm. (D) Percentage of cells with nucleolar luciferase foci after heat stress and recovery ± CHIP overexpression. Mean ± SD from n = 2 experiments. Two-way ANOVA with Tukey’s test. (E) Quantification of cells with DRiPs foci after CHIP overexpression ± HSP70 inhibition. Mean ± SD from n = 3 experiments. Two-way ANOVA with Tukey’s test. (F) Percentage of cells with nucleolar luciferase foci during recovery in the presence of CHIP WT, H260Q, or K30A. Mean ± SD from n = 3 experiments. Two-way ANOVA with Tukey’s test. (G) Pearson’s correlation coefficient between nucleolar luciferase and HSP70 ± VER. Mean ± SD from n = 3 experiments. Welch’s t test. (H) Pearson’s correlation coefficient between nucleolar luciferase and NPM1 under the conditions shown in (G). Mean ± SD from n = 3 experiments. Welch’s t test. See also Figure S2

To evaluate how HSP70 activity contributes to Ub handling, we modulated the chaperone activity with VER and an activator (SW02, a small-molecule allosteric activator that stabilises the ADP-bound state and prolongs HSP70-substrate interactions)^32,33^. Although absolute nucleolar Ub intensity was variable (Fig. S2B1), VER consistently decreased the nucleolar/nucleoplasmic Ub ratio (Fig. S2B2), indicating that HSP70 activity contributes to maintaining Ub substrate flux beyond its classical folding functions.

To directly probe CHIP role in sequestration, we employed a thermolabile nuclear luciferase reporter^4^. Luciferase accumulated in nucleoli during heat and dispersed upon recovery, colocalising with CHIP in foci (Fig. 2C). Quantification showed that CHIP expression alone, but not HSP70, significantly increased nucleolar sequestration of luciferase, with no additive effect of co-expression, indicating CHIP is the dominant driver (Fig. 2D).

We next examined nascent misfolded proteins using O-propargyl-puromycin (OPP) labelling combined with click chemistry to visualise defective ribosomal products (DRiPs), i.e. prematurely terminated or misfolded nascent chains. Both wild-type CHIP and the catalytically inactive CHIP^H260Q^ variant promoted DRiPs accumulation in nucleoli, whereas the HSP70-binding-defective CHIP^K30A^ mutant^34,35^ failed to do so (Fig. 2E, S2C). These findings suggest that CHIP functions primarily as an HSP70 adaptor, rather than as an E3 ligase, to expand PQC capacity.

We then asked whether enhanced sequestration alters clearance. In luciferase reporter cells, both CHIP and CHIP^H260Q^ delayed foci dissolution during recovery, with CHIP^H260Q^ causing persistent nucleolar retention up to 6 h (Fig. 2F). By contrast, CHIP^K30A^ had minimal effect, underscoring that CHIP requires HSP70 binding to efficiently modulate clearance dynamics, whereas its E3 ligase activity (H260Q) is critical for substrate release. To generalise this result, we turned to chromobox protein homolog 2 (CBX2), a misfolding-prone, HSP70-dependent client previously shown to accumulate in nucleoli upon heat stress^30^. Both CHIP and CHIP^H260Q^ delayed CBX2 clearance (Fig. S2D). Pharmacological inhibition of HSP70 with VER mirrored CHIP overexpression by slowing clearance, and combined CHIP overexpression with VER exacerbated this defect (Fig. S2E). Together, this suggests that CHIP maintains nucleoli in a proteostasis-active state by modulating HSP70 substrate processing.

Finally, to dissect the architectural basis, we analysed spatial relationships among luciferase, HSP70 and NPM1. CHIP reduced luciferase-HSP70 proximity while increasing luciferase-NPM1 colocalisation (Fig. 2G-H). Notably, VER treatment reproduced these effects in control cells, consistent with CHIP overabundance driving a chaperone-inhibited state. Quadruple staining confirmed co-existence of CHIP, HSP70, NPM1, and luciferase in nucleolar foci (Fig. S2F1-F3), supporting a model in which CHIP alters the local stoichiometry of the PQC triad, thereby shifting clients into an NPM1-rich microenvironment. In summary, CHIP drives HSP70-dependent nucleolar sequestration of misfolded proteins, expanding storage but delaying clearance and recovery.

### CHIP-driven Nucleolar Remodeling Couples Protein Storage with Inhibition and Reorganisation of Ribosome Biogenesis During Stress Recovery

Having shown that CHIP promotes nucleolar storage of misfolded proteins, we next asked whether its sustained retention interferes with recovery processes. Perturbation of Ub-dependent clearance confirmed its persistance: proteasome inhibition with MG132 or blocking deubiquitinating enzymes with PR-619 delayed CHIP removal, whereas inactivation of cullin-RING ligases by MLN4924^36^had little effect (Fig. 3A1). Under these conditions, nucleoli accumulated Annexin A5 (ANXA5), a stress-inducible annexin previously identified in nucleolar proteomes under heat shock^4^, along with ubiquitinated proteins (Fig. 3A2-3). Notably, these nucleoli failed to restore Bystin, a factor required for rRNA processing^37^ (Fig. 3A4), suggesting that sustained PQC engagement compromises ribosome biogenesis reactivation.

**Figure 3.**
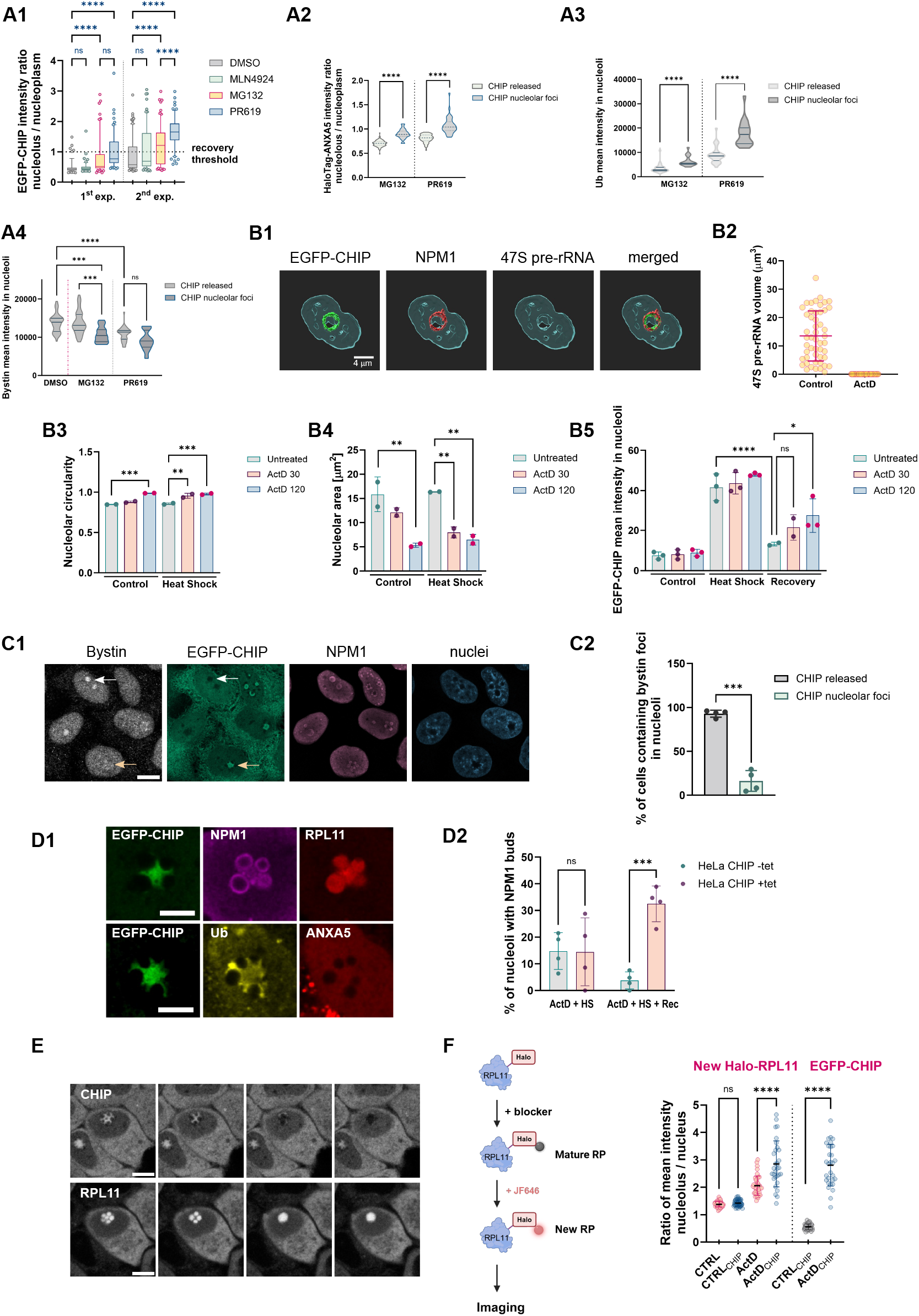
CHIP-driven Nucleolar Remodeling Links Protein Storage to Ribosome Biogenesis. (A) Ubiquitin and proteasome activity regulate CHIP and substrate clearance. (A1) CHIP redistribution classified by nucleolar/nucleoplasmic intensity ratio. (A2-A4) Effects of MG132 and PR619 on HaloTag-ANXA5 localisation and on nucleolar ubiquitin and Bystin levels. Box-and-whisker plots show median and 5^th^-95^th^ percentiles (CHIP); violin plots show median and quartiles (others). n = 15-99 nucleoli per condition; 2 experiments. Statistical tests: Welch’s t-test or ANOVA with post-hoc Tukey’s test. (B) Inhibition of rRNA synthesis alters nucleolar morphology and CHIP clearance. (B1-B2) FISH for 47S pre-rRNA and quantification of nucleolar volume upon ActD. Scale bar, 4 µm. (B3-B4) Morphometric analysis of nucleolar circularity and area. (B5) CHIP nucleolar intensity. Mean ± SD, n = 2–3 experiments; ANOVA with Dunnett’s test. (C) CHIP and Bystin show reciprocal dynamics. (C1) Confocal images of CHIP/Bystin redistribution during recovery. White arrows, Bystin-enriched nucleoli; orange arrows, delayed CHIP clearance. Scale bar, 10 µm. (C2) Quantification of nucleolar Bystin enrichment. Mean ± SD, n = 4; Welch’s t-test. (D) CHIP induces subcompartmentalised nucleolar morphology. (D1) Confocal images of NPM1-positive nucleolar protrusions containing HaloTag-ANXA5 or RPL11. Scale bar, 5 µm. (D2) Quantification of “budded” nucleoli. Mean ± SD, n = 4; two-way ANOVA with Sidak’s test. (E) Time-lapse imaging of CHIP-to-RPL11 exchange in budded nucleoli. Four frames shown. Scale bar, 5 µm. See also Video S1. (F) CHIP traps nascent RPL11 in nucleoli. Schematic of HaloTag-RPL11 pulse labelling. Quantification of nucleolar/nuclear intensity ratios for nascent RPL11 and EGFP-CHIP. (CTRL, ActD: HeLa CHIP (-tet); CTRL_CHIP_, ActD_CHIP_: HeLa CHIP (+tet). Mean ± SD, n = 30-62 nucleoli per condition; 3 experiments; Welch’s t-test. See also Figure S3

To probe this interplay more directly, we combined heat shock with Actinomycin D (ActD), an RNA polymerase I inhibitor that suppresses pre-rRNA synthesis and induces nucleolar caps^38,39^. ActD inhibited rRNA transcription (Fig. 3B1-2), remodelled nucleoli into smaller circular structures with FBL-rich caps (Fig. 3B3-4; Fig. S3A1-2), and significantly delayed CHIP clearance after stress (Fig. 3B5, S3B1-2), thereby revealing a checkpoint coupling ribosome synthesis to proteostasis resolution. ActD also altered CHIP distribution, concentrating it with NPM1 in ring-like structures (Fig. 3B1, S3A1, S3B1). This setting allowed us to capture transition events between proteostasis and biosynthesis. During ActD and heat shock co-treatment, Bystin partially effluxed to the nucleoplasm, resulting in a strong reduction of its nucleolar levels (Fig. S3C). Strikingly, during recovery, Bystin gradually re-accumulated in nucleoli despite continued ActD treatment, coinciding with CHIP release. This reciprocal CHIP-Bystin exchange (Fig. 3C1-2) highlights a regulated switch from PQC to ribosome biogenesis.

We also identified distinct morphological correlates of this transition. In CHIP-overexpressing cells, recovering nucleoli frequently adopted a ‘flowering’ architecture (Fig. 3D1-2), characterised by CHIP and ubiquitin concentrated in the core, ribosomal protein RPL11 arranged in peripheral ‘petals,’ and NPM1 forming an outer ring. Time-lapse imaging revealed that these structures subsequently coalesced into spherical nucleoli as CHIP was replaced by RPL11 (Fig. 3E; Video S1), recapitulating the exchange previously observed between CHIP and Bystin. Finally, to assess whether CHIP affects ribosome biogenesis directly, we tracked orphan ribosomal proteins using nascently labelled Halo-RPL11. In CHIP-overexpressing cells, RPL11 persisted longer in nucleoli, consistent with delayed incorporation into ribosomal subunits or impaired recycling (Fig. 3F). In summary, CHIP-driven sequestration antagonises ribosome biogenesis by reorganising nucleoli and prolonging the proteostasis-active state.

### CHIP Accumulation in Nucleoli Links Proteostasis to Transcriptomic Remodeling of Ribosome Biogenesis

Establishing that CHIP accumulates in nucleoli under heat stress and prolongs the proteostasis-engaged state, we next investigated whether endogenous CHIP exhibits similar localisation dynamics and influences the transcriptional landscape of nucleolar function. To this end, we fractionated HeLa Flp cells into cytoplasmic, nucleoplasmic, and nucleolar fractions under control and heat shock conditions. Fractionation of HeLa Flp cells into cytoplasmic, nucleoplasmic, and nucleolar fractions confirmed that both CHIP and HSP70 are enriched in the nucleolar fraction following heat shock (Fig. 4A). Immunofluorescence in HeLa Flp and MCF7 cells similarly revealed endogenous CHIP relocalisation to nucleoli, validated by quantitative intensity analysis (Fig. 4B, S4A1-2).

**Figure 4.**
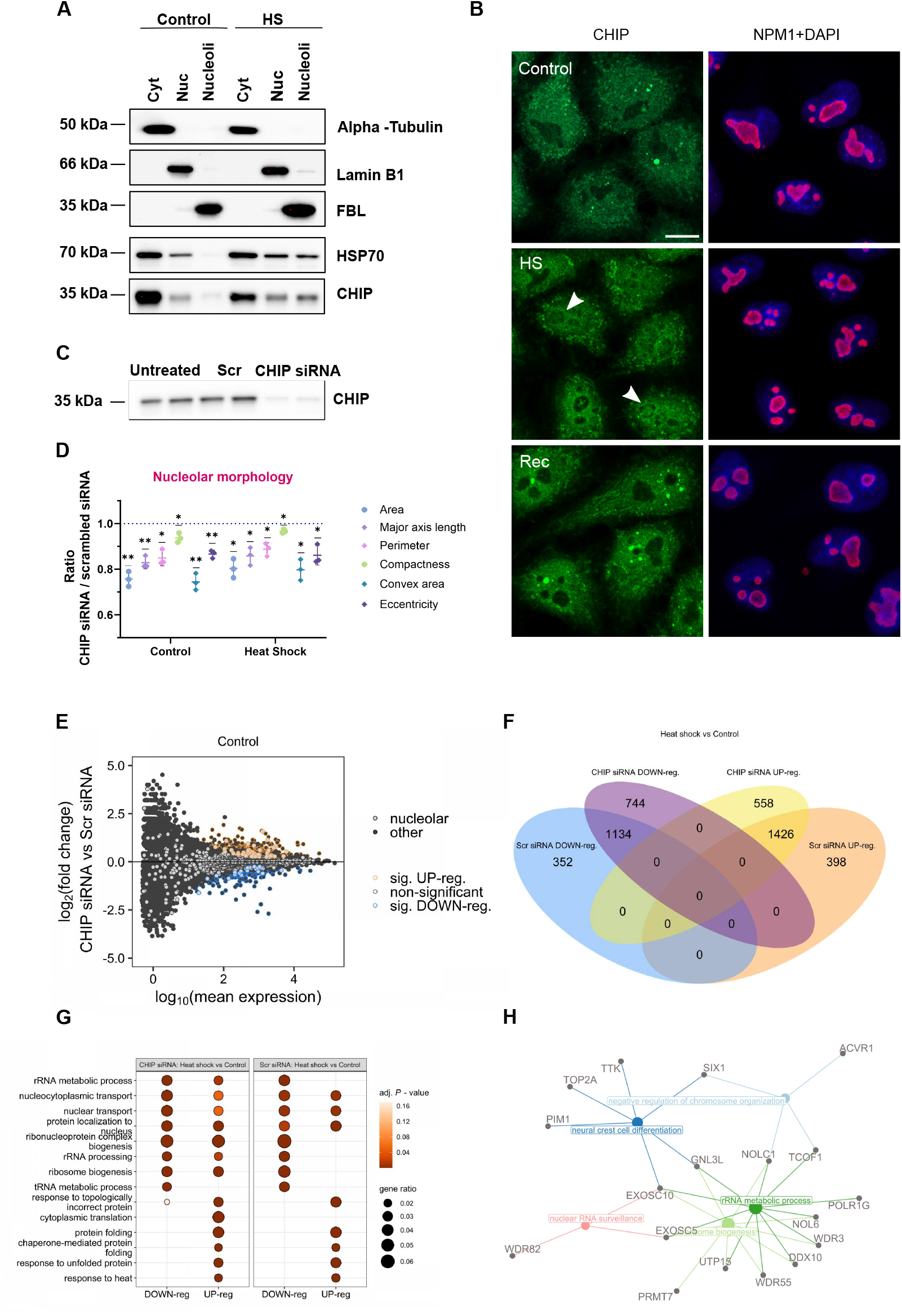
CHIP Links Nucleolar Proteostasis to Transcriptomic Remodeling of Ribosome Biogenesis. (A) Western blot of HeLa Flp cells fractionated into cytoplasmic (Cyt), nucleoplasmic (Nuc), and nucleolar fractions, showing CHIP and HSP70 enrichment in nucleoli after heat shock. Fraction purity was assessed by α-tubulin (cytoplasm), Lamin B1 (nucleoplasm), and FBL (nucleoli). (B) Endogenous CHIP relocalises to nucleoli upon heat shock. Immunofluorescence of CHIP (green) and NPM1 (red) in control, heat-stressed (HS), and recovered (Rec) cells. Scale bar, 10 µm. (C) Western blot validation of CHIP knockdown by siRNA. Scrambled siRNA (Scr) served as control. (D) CHIP depletion alters nucleolar morphology. Quantification of morphological parameters under basal and heat-shock conditions. Mean ± SD from n = 3 experiments; one-sample t-test. (E) RNA-seq analysis of siCHIP versus siScr cells under basal conditions. MA plot showing global differential expression. Genes encoding nucleolar proteins are highlighted in light grey, significantly downregulated genes in blue, and upregulated genes in orange (adj. p < 0.05, Wald test with Benjamini-Hochberg correction). (F) Venn diagram of genes deregulated by heat shock in control versus CHIP-depleted cells. (G) GO enrichment analysis of biological processes affected in CHIP-depleted cells under heat shock. Dot size indicates the gene ratio, colour indicates adjusted p-value. (H) Gene-pathway network analysis of selected deregulated genes in CHIP-depleted cells under basal conditions. See also Figure S4

To probe functional consequences, we depleted CHIP by siRNA (Fig. 4C, S4B). Even under basal conditions and during heat shock, CHIP knockdown induced subtle nucleolar defects, with nucleoli appearing smaller and more rounded, reminiscent of ActD treatment (Fig. 4D), suggesting a role for CHIP in maintaining nucleolar integrity and stress adaptability. RNA-seq comparing siCHIP and Scr-siRNA cells under basal, heat shock, and recovery conditions revealed widespread transcriptional changes. Even without stress, CHIP knockdown altered the expression of hundreds of genes, with nucleolar factors significantly enriched among the deregulated transcripts (Fig. 4E, S4C, Table S1). Upon heat shock, the divergence between CHIP-depleted and control cells became more pronounced. CHIP knockdown led to a greater number of uniquely regulated genes, with 558 vs. 398 uniquely upregulated and 744 vs. 352 uniquely downregulated compared to controls (Fig. 4F, Table S2). Gene Ontology (GO) analysis revealed a significant enrichment of rRNA metabolic processes, rRNA processing, and ribosome biogenesis among the upregulated transcripts (Fig. 4G, Table S3), indicating that CHIP normally regulates these pathways under stress. Network visualisation of CHIP-depleted cells revealed that deregulated transcripts are interconnected, especially within rRNA metabolism and ribosome biogenesis pathways (Fig. 4H). Among these, genes such as NOL6 (SSU processome component), WDR55 (rRNA transcription and processing), NOLC1 (nucleolar scaffold regulating Pol I and snoRNP trafficking), UTP15 (SSU processome component), WDR18 (PELP1 complex factor in 60S maturation), and GNL3L (nucleolar GTPase in rRNA processing and cell cycle control) stand out as candidates for future mechanistic dissection of how nucleolar PQC transitions are coupled to transcriptional control of biosynthetic recovery^40–44^.

### NOL6 and WDR55 Differentially Modulate CHIP-dependent Nucleolar Storage Responses

To functionally validate CHIP-sensitive transcripts from RNA-seq, we performed an siRNA screen against nucleolar genes annotated for rRNA processing and structure. Using HeLa CHIP cells, we monitored CHIP localisation during heat shock and recovery. Among the candidates, NOL6 depletion reduced CHIP sequestration during stress, whereas WDR55 or GNL3L depletion impaired CHIP clearance during recovery (Fig. 5A). Based on the robustness and opposing nature of these phenotypes, we prioritised NOL6 and WDR55 for further study.

**Figure 5.**
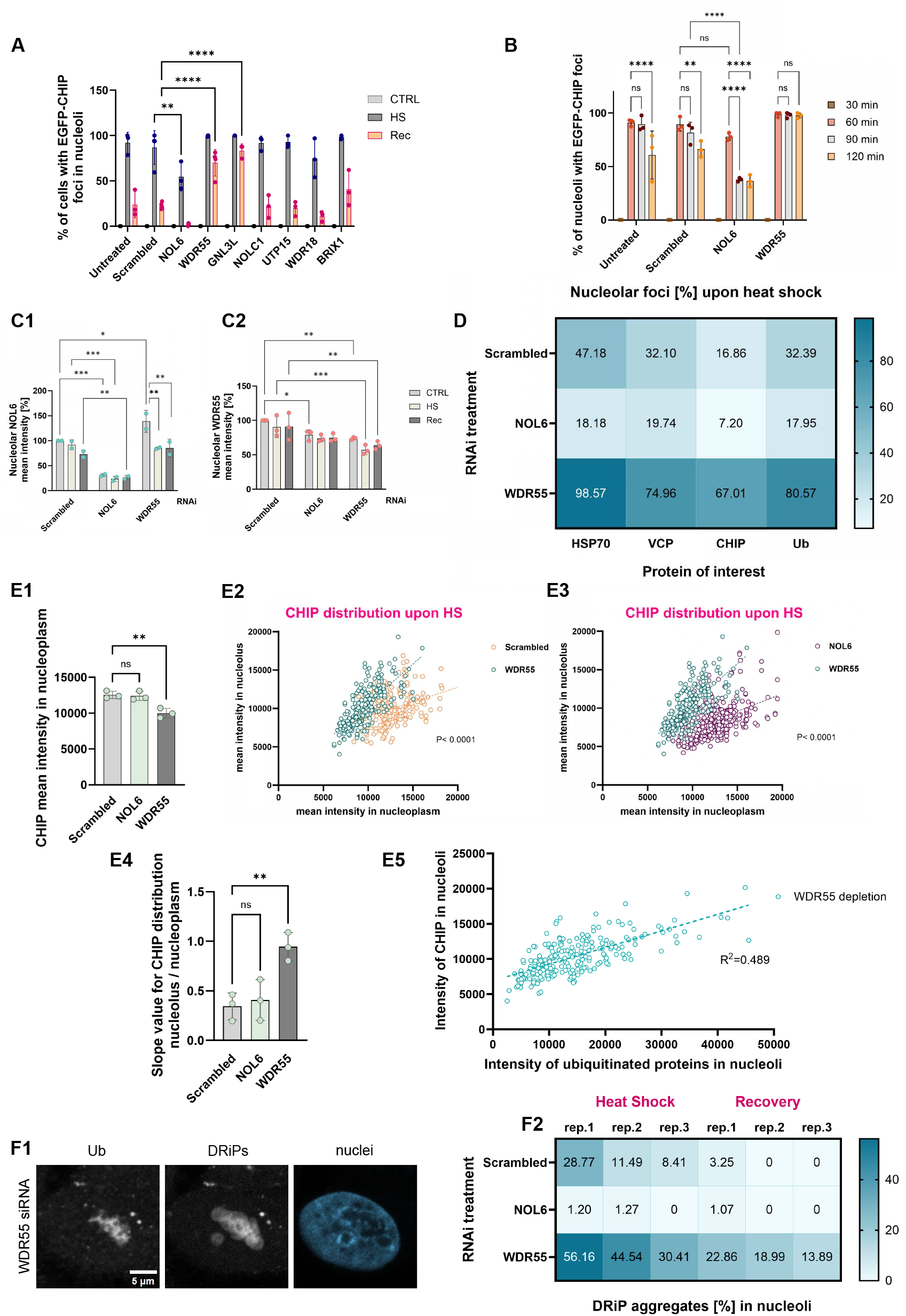
Intrinsic nucleolar factors NOL6 and WDR55 tune CHIP-dependent storage capacity. (A) Quantification of EGFP-CHIP sequestration in nucleoli upon depletion of various nucleolar proteins during heat shock and recovery (mean ± SD, n = 3). Two-way ANOVA with Dunnett’s test. (B) NOL6 knockdown accelerates CHIP release from nucleoli during stress (mean ± SD, n = 3, two-way ANOVA with Tukey’s test). (C) Knockdown validation for NOL6 (C1) and WDR55 (C2) (mean ± SD, n = 2-3). (D) Recruitment of PQC factors to nucleoli is reduced by NOL6 depletion but exaggerated by WDR55 depletion. Percentage of nucleolar foci (intensity ratio nucleolus/nucleoplasm > 1), (n = 3). (E) WDR55 loss enhances CHIP-ubiquitin coupling. (E1-E4) Quantification of CHIP intensity and distribution between nucleolus and nucleoplasm after heat shock across RNAi treatments with nonlinear regression and slope analysis (mean ± SD, n = 3). (E5) Correlation between CHIP and ubiquitin recruitment in WDR55-depleted cells (Pearson r = 0.6693, R2=0.489, P < 0.0001). (F) WDR55 depletion causes persistence of DRiP foci. (F1) Representative confocal images of DRiPs and ubiquitin in nucleoli after OPP treatment (scale bar, 5 µm). (F2) Quantification of nucleoli with DRiP aggregates across RNAi treatments (n = 3). See also Figure S5

Although their roles in human nucleolar stress responses remain poorly defined, orthologue-based studies highlight broader relevance. NOL6 is a conserved RNA-binding protein and a core component of the small subunit (SSU) processome, required for the early cleavage of 47S pre-rRNA during 40S biogenesis^45,46^. In mammalian cells, NOL6 has been linked to proliferation and cell cycle control, whereas in *C. elegans* its orthologue modulates innate immunity, ER stress adaptation, and longevity^47–49^. WDR55, by contrast, is a WD40-repeat protein localising to the DFC, partially co-distributed with FBL^50^. In plants, it functions as a substrate adaptor for cullin 4 ubiquitin ligases^51^, suggesting possible links to proteostasis.

Cell imaging showed that CHIP accumulated in nucleoli with similar kinetics across all conditions, peaking ~1 h into heat shock. However, in NOL6-depleted cells CHIP exited nucleoli prematurely, consistent with compromised storage capacity and accelerated recovery. In contrast, WDR55 knockdown caused CHIP to persist, locking cells in a storage-active state (Fig. 5B). Immunofluorescence confirmed knockdown efficiencies (~75% for NOL6, ~25% for WDR55), with both proteins enriched in nucleoli (Fig. 5C, S5A). In addition, WDR55 decreased level altered its redistribution across the nuclear region during heat shock, as compared to scrambled and NOL6-depleted cells (Fig. S5B).

To assess the broader PQC architecture, we quantified HSP70, CHIP, Ub conjugates, and valosin-containing protein (VCP/p97), an AAA+ ATPase central to Ub-dependent protein degradation across the cytoplasm, nucleus, and ER, and previously reported to localise to nucleoli^52^. NOL6 knockdown reduced nucleolar recruitment of all markers, whereas WDR55 depletion exaggerated sequestration (Fig. 5D, S5C). Microscopy further showed reduced nucleoplasmic CHIP in WDR55-depleted cells, yet strong nucleolar accumulation and increased Ub signal (Fig. 5E). This suggests that WDR55 deficiency stabilises a hyper-sequestration state by impairing recovery-associated CHIP release. Importantly, defective resolution extended to DRiPs: persistent DRiPs foci were observed in WDR55-deficient nucleoli after heat shock (Fig. 5F), despite normal translation rates (Fig. S5D), indicating that delayed clearance was not due to increased substrate production. In summary, NOL6 facilitates nucleolar PQC engagement, whereas WDR55 enables exit from the storage state, defining opposing regulators that tune CHIP-dependent stress responses.

### Integration of Nucleolar PQC With Cellular Proteostasis via VCP and ISR

Our data so far indicate that nucleolar proteostasis is an actively remodelled state, shaped by CHIP and regulators such as NOL6 and WDR55. To test whether this compartment functions in isolation or as part of a coordinated cellular network, we examined the interplay between nucleolar PQC and global proteostasis pathways.

Silencing of WDR55 markedly increased nucleolar accumulation of active proteasomes during heat shock (Fig. 6A1-A2), consistent with nucleoli entering a storage-dominated PQC state. We next focused on VCP, given its recruitment to nucleoli during stress. VCP inhibition with CB5083 induced cytoplasmic foci enriched in VCP and Ub conjugates, while simultaneously reducing nuclear Ub and suppressing stress-induced recruitment of both VCP and Ub into nucleoli. In contrast, treatment with SMER28, a small-molecule activator of VCP that also promotes autophagy, had little effect, suggesting that VCP activity is not rate-limiting in this context (Fig. 6B1-B5). These findings indicate that VCP is required for nucleolar entry and inter-compartmental flux of ubiquitinated proteins, thereby sustaining PQC plasticity during stress. Consistent with this role, CB5083 disrupted nuclear Ub-CHIP foci, also situated in close proximity to nucleoli, in WDR55-depleted cells (Fig. 6C1), and reduced overall nucleolar Ub burden despite largely preserved CHIP localisation (Fig. 6C2, S6A). Notably, VCP inhibition also impaired WDR55 recruitment to nucleoli, rerouting it into cytoplasmic foci (Fig. S6B), suggesting that VCP activity stabilises not only client flux but also the localisation of regulatory factors.

**Figure 6.**
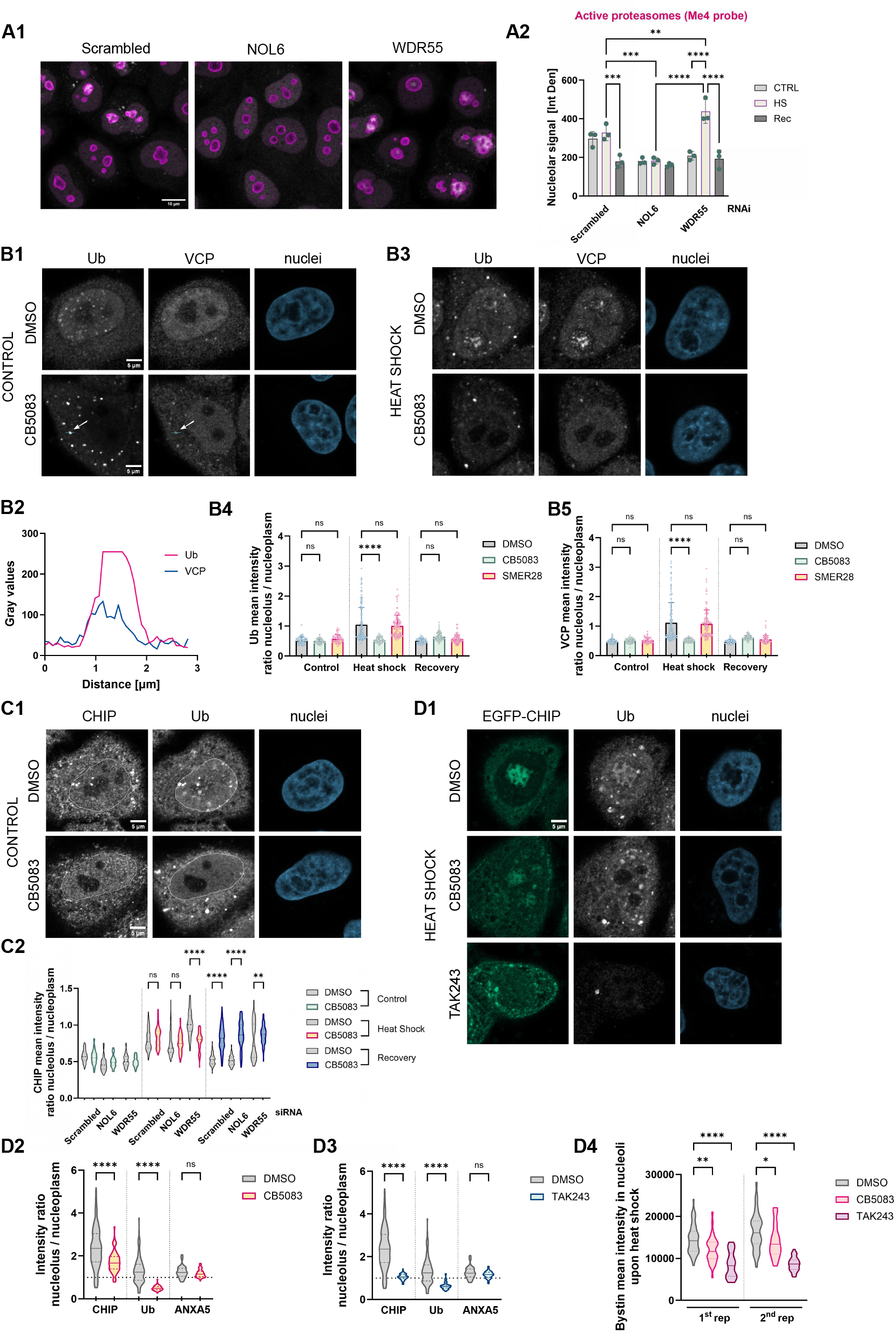
VCP and ISR Integrate Nucleolar PQC with Cellular Proteostasis. (A) WDR55 depletion enhances nucleolar proteasome activity during heat shock. (A1) Confocal images of HeLa Flp cells after heat shock labelled with the proteasome activity probe Me4 after the indicated RNAi treatments (NPM1, magenta; Me4, grey). Scale bar, 10 µm. (A2) Quantification of nucleolar Me4 intensity in basal, heat-shock, and recovery conditions. Mean ± SD from n = 3 experiments; two-way ANOVA with Tukey’s test. (B) VCP inhibition suppresses nucleolar recruitment of Ub conjugates and VCP while inducing cytoplasmic foci. (B1, B3) Confocal images of Ub and VCP distribution in HeLa Flp cells treated with DMSO or CB5083 under basal and heat shock conditions. White arrows mark CB5083-induced cytoplasmic foci enriched in Ub and VCP. Scale bars, 5 µm. (B2) Intensity profiles of Ub and VCP signals across the cytoplasmic focus shown in (B1). (B4-B5) Quantification of nucleolar/nucleoplasmic Ub and VCP ratios. Individual values shown from one experiment; mean ± SD. Data representative of n = 3 experiments. (C) VCP inhibition disrupts CHIP-Ub trafficking and delays CHIP clearance during recovery. (C1) Confocal images of HeLa Flp cells stained for CHIP and Ub conjugates. Dashed lines mark nuclei. Scale bar, 5 µm. (C2) Quantification of CHIP nucleolar/nucleoplasmic ratios in control, heat shock, and recovery ± CB5083. Violin plots show median and quartiles (n = 357 [CTRL], 790 [HS], 541 [Rec] nucleoli). One-way ANOVA with Sidak’s test. (D) Inhibition of Ub activation phenocopies VCP inhibition. (D1) Confocal images of HeLa CHIP cells pretreated with CB5083 or TAK243 before heat shock, stained for Ub conjugates. Scale bar, 5 µm. (D2-D3) Quantification of nucleolar/nucleoplasmic EGFP-CHIP, Ub, and HaloTag-ANXA5 ratios in heat-shocked cells. Violin plots show median and quartiles (n = 110 [DMSO], 79 [CB5083], 75 [TAK243] nucleoli). Data pooled from n = 2 experiments; one-way ANOVA with Tukey’s test. (D4) Bystin nucleolar levels in the same conditions. Violin plots show median and quartiles (n = 105 [DMSO], 75 [CB5083], 68 [TAK243] nucleoli across 2 repeats). One-way ANOVA with Tukey’s test. See also Figure S6

To further dissect the involvement of the UPS, we compared CB5083 with the Ub-activating enzyme inhibitor TAK243 in HeLa CHIP cells. Both inhibitors suppressed nucleolar Ub accumulation during heat shock (Fig. 6D1-D3). TAK243, however, more strongly reduced CHIP recruitment, whereas CB5083 primarily altered its nucleoplasmic distribution (Fig. 6D1-D2, S6C). Neither treatment prevented ANXA5 accumulation (Fig. 6D2-D3), but both reduced nucleolar Bystin levels (Fig. 6D4), indicating disruption of the PQC-biosynthesis balance. Addition of CB5083 specifically during recovery did not block CHIP clearance (Fig. S6D1), but selectively reduced Bystin in CHIP-positive nucleoli (Fig. S6D2). Moreover, under both inhibitor conditions, Bystin relocalised into cytoplasmic foci overlapping with the stress granule marker FXR1 (Fig. S6E), suggesting that when nucleolar recovery fails, biosynthetic factors are rerouted to stress granules as auxiliary proteostasis sites.

Finally, we tested whether ISR, a conserved eIF2α-dependent translational checkpoint that also represses rRNA processing, coordinates this transition. Inhibition of ISR with ISRIB reversed the NOL6 knockdown phenotype: instead of premature CHIP release, cells retained CHIP in nucleoli, resembling WDR55-deficient conditions (Fig. S6F top). Continued ISRIB treatment further impaired recovery and sustaining EGFP-CHIP in nucleoli across all RNAi backgrounds, resulting in the formation of cytoplasmic foci (Fig. S6F bottom). These findings support a role for ISR in promoting exit from the CHIP-engaged state, thereby creating conditions that permit the resumption of ribosome biogenesis. Together with VCP, ISR integrates nucleolar PQC with cellular proteostasis, balancing storage and recovery

### Magnitude of Nucleolar Reorganisation Reflects the Balance Between PQC Engagement and Ribosome Biogenesis

Building on the role of VCP and ISR as global integrators of nucleolar PQC (Fig. 6), we next asked how nucleolar-intrinsic factors such as NOL6 and WDR55 shape adaptation to stress. Immediately after heat shock, NOL6 knockdown increased both Bystin and ubiquitin signal in CHIP-rich nucleoli (Fig. 7A), correlating with faster recovery. These nucleoli also adopted a distinctive rounded morphology compared with scrambled controls or WDR55-depleted cells (Fig. 7B1-B2), suggesting that NOL6 normally stabilises rRNA metabolism and influences nucleolar susceptibility to proteotoxic stress.

**Figure 7.**
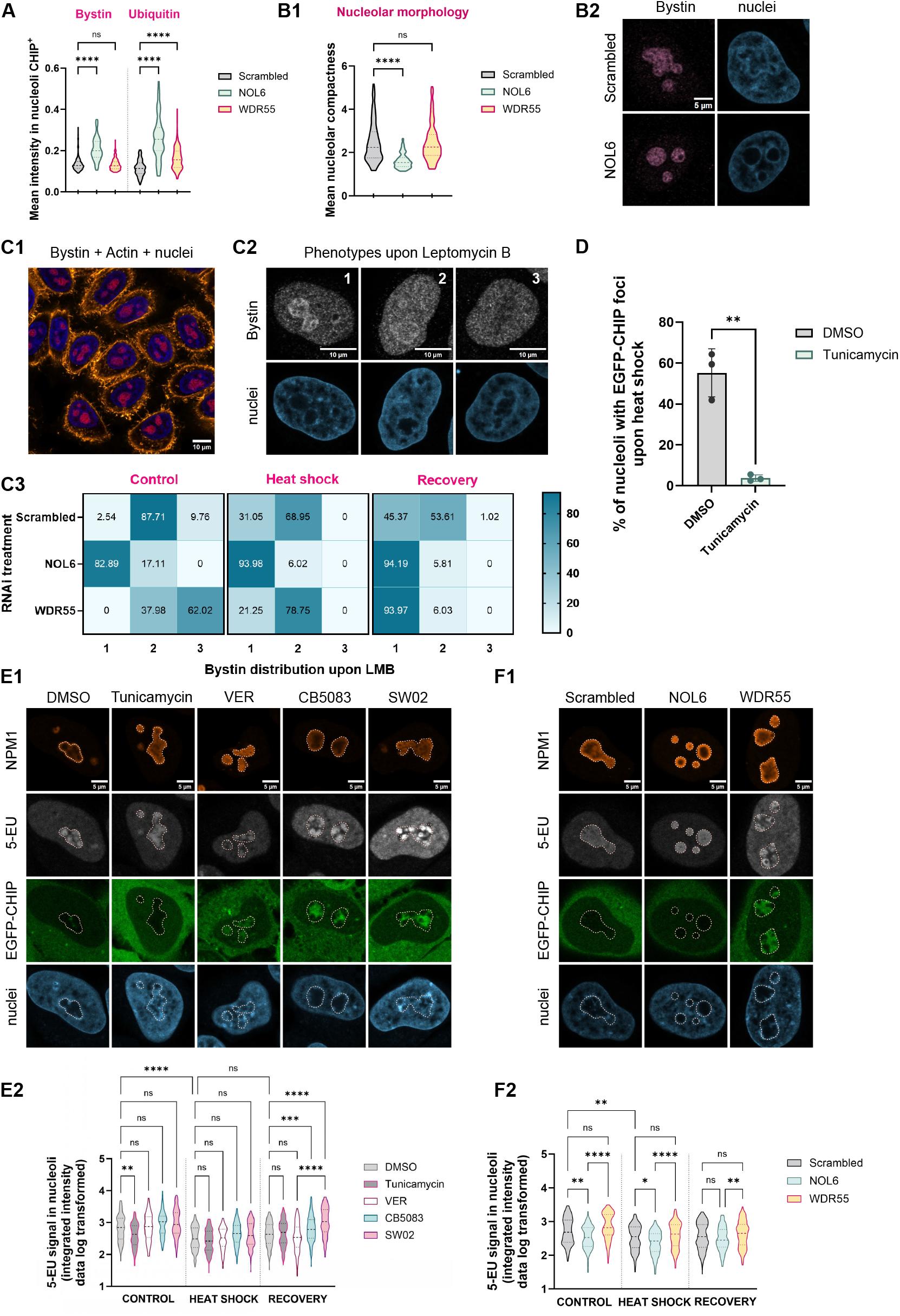
Nucleolar Reorganisation Balances PQC Engagement with Ribosome Biogenesis. (A) NOL6 depletion increases Bystin and Ub accumulation in CHIP-rich nucleoli. Violin plots show median and quartiles (n = 157 [siScr], 68 [siNOL6], 153 [siWDR55] nucleoli, one repeat; experiments repeated n = 2 [Ub] or 3 [Bystin]. One-way ANOVA with Tukey’s test. (B) NOL6 depletion enhances nucleolar compactness during heat stress. (B1) Violin plots of nucleolar circularity (n = 193 [siScr], 257 [siNOL6], 159 [siWDR55] nucleoli, one repeat; n = 3 experiments. One-way ANOVA with Tukey’s test. (B2) Representative confocal images of nucleoli stained for Bystin. Scale bar, 5 µm. (C) Leptomycin B (LMB) alters Bystin dynamics differently in NOL6- and WDR55-depleted cells. (C1) Confocal image of HeLa Flp cells in basal conditions (Bystin, red; actin, orange). (C2) Representative images of phenotypic classes based on Bystin distribution after LMB treatment. Scale bars, 10 µm. (C3) Frequency of phenotypes under indicated RNAi conditions during basal, heat-shock, and recovery states. Mean from n = 2 experiments. (D) ER stress reduces CHIP recruitment to nucleoli. Quantification of nucleoli with EGFP-CHIP foci in heat-shocked cells ± tunicamycin pretreatment. Mean ± SD from n = 3 experiments. Unpaired t-test. (E) SW02 and CB5083 increase rRNA synthesis during recovery. (E1) Confocal images of HeLa CHIP cells pulsed with 5-EU to label nascent rRNA in nucleoli. Dashed lines mark nucleoli. Scale bar, 5 µm. (E2) Quantification of 5-EU intensity under basal, heat shock, and recovery conditions. Violin plots show median and quartiles (n = 134-341 nucleoli/condition, n = 2 experiments). Kruskal-Wallis with Dunn’s test. (F) NOL6 depletion decreases rRNA production. (F1) Confocal images of HeLa CHIP cells during recovery after siRNA knockdown of NOL6 or WDR55, pulsed with 5-EU. Dashed lines mark nucleoli. Scale bars, 5 µm. (F2) Quantification of 5-EU intensity across conditions. Violin plots show median and quartiles (n = 123-232 nucleoli/condition, n = 2 experiments). Kruskal-Wallis with Dunn’s test. See also Figure S7

To determine whether altered Bystin abundance reflected accelerated ribosome assembly or disrupted Bystin dynamics, we used leptomycin B (LMB), an inhibitor of nucleocytoplasmic transport that unmasks 40S assembly defects by trapping Bystin in nucleoli^53^ (Fig. S7A). Three reproducible phenotypes were distinguished (Fig. 7C1-2): (1) strong nucleolar Bystin retention, (2) reduced nucleolar Bystin comparable to nucleoplasmic levels, and (3) near-complete release, giving the appearance of nucleolar emptying. Scrambled controls predominantly adopted phenotype 2, consistent with normal 40S assembly (Fig. 7C3). NOL6 depletion shifted nucleoli toward phenotype 1 even under basal conditions, indicating impaired dynamics and slowed ribosome assembly. By contrast, WDR55 depletion biased toward phenotype 3, suggesting excessive Bystin efflux. Heat shock partially normalised WDR55-deficient nucleoli toward phenotype 2, but NOL6-deficient nucleoli remained locked in phenotype 1. During recovery, control cells remained in phenotype 2, whereas WDR55-deficient nucleoli shifted to phenotype 1 and both knockdowns were maintained in aberrant states. Thus, although both factors modulate CHIP-dependent PQC, NOL6 depletion perturbs early stages of rRNA metabolism and restricts storage capacity, whereas WDR55 depletion enforces a storage-biased state that prevents re-entry into ribosome biogenesis. These divergent effects extended to other PQC substrates. In NOL6-depleted cells, DRiP-positive foci were markedly reduced both under basal and heat-shock conditions (Fig. S7B1-B2), suggesting that aberrant nascent proteins are channelled away from nucleolar storage, consistent with the faster recovery phenotype. By contrast, scrambled and WDR55-depleted cells retained robust DRiPs accumulation together with ubiquitin in nucleoli after heat shock (Fig. S7B2), indicating a sustained sequestration mode in the absence of WDR55.

We next tested whether stress originating outside the nucleolus rewires its PQC function. Pretreatment with tunicamycin, a classical ER stressor that blocks N-linked glycosylation, significantly reduced CHIP accumulation in nucleoli during heat shock (Fig. 7D). This effect likely reflects redistribution of PQC resources toward ER stress resolution, although metabolic suppression by tunicamycin may also contribute. Together, these data highlight that nucleolar PQC is dynamically coupled to cytoplasmic and ER stress responses, with CHIP partitioning reflecting global prioritisation of proteostasis resources. To further link proteostasis with ribosome output, we monitored rRNA synthesis using 47S pre-rRNA FISH (Fig. S7C) and 5-EU incorporation (Fig. 7E-F). Control cells resumed rRNA production within 1 h of recovery, whereas CHIP-overexpressing cells remained suppressed, resembling VER-treated conditions (Fig. S7C). Quantification confirmed that heat shock reduced rRNA synthesis across all treatments (Fig. 7E2, 7F2). NOL6 depletion further suppressed rRNA output, consistent with impaired early steps of ribosome assembly as also indicated by LMB sensitivity (Fig. 7C3, 7F2). Tunicamycin likewise decreased rRNA production, most likely reflecting global metabolic suppression and redistribution of PQC resources (Fig. 7E2). In contrast, SW02 (HSP70 activator) and CB5083 (VCP inhibitor) paradoxically enhanced rRNA synthesis during recovery, even though CHIP clearance remained incomplete (Fig. 7E1–E2, S7D). This suggests that HSP70 and VCP influence rRNA transcription through mechanisms uncoupled from PQC resolution.

Finally, to capture structural correlates of these transcriptional states, we classified nucleoli into three phenotypes based on rRNA distribution (Fig. S7E): free-flow (continuous rRNA flux), sand-like (rRNA sealed within CHIP-enriched foci, most prominent under VER) and peripheral (rRNA displaced toward the rim). NOL6 knockdown and tunicamycin both enriched the free-flow phenotype even during stress, while VER abolished it entirely (Fig. 7F1, Fig. SF1-F2). By contrast, SW02 increased the frequency of peripheral nucleoli, a morphology we hypothesised to represent an adaptive state accommodating elevated rRNA flux while PQC remains engaged. Fisher’s exact test confirmed that VER-treated nucleoli were significantly less likely to adopt the peripheral phenotype compared with DMSO, CB5083, or SW02 conditions (Fig. S7F3). Together, these findings reveal that nucleolar reorganisation proceeds through distinct morphological states corresponding to functional fates: NOL6 depletion and tunicamycin bias toward premature rRNA flux, WDR55 depletion locks nucleoli in storage, VER imposes a sealed state refractory to recovery, and SW02 promotes a peripheral phenotype that may represent an adaptive intermediate.

## Discussion

Our findings expand the emerging concept of the nucleolus as a proteostasis hub that not only produces ribosomes but also dynamically reorganises to buffer proteotoxic stress. Previous studies have shown that misfolded proteins, including defective ribosomal products, luciferase, and polycomb group proteins, accumulate in nucleoli during heat shock or proteasome inhibition and are subsequently cleared in recovery through proteasome activity^4,5,54^. This positioned the nucleolus as a transient storage compartment. Here, we refine this model by showing that the process is not passive: CHIP and VCP actively govern the flux of substrates, while nucleolar biogenesis factors such as NOL6 and WDR55 modulate the amplitude and kinetics of storage and recovery. Importantly, we demonstrate that ubiquitination within nucleoli is highly structured and functions as a regulatory signal, committing proteins to sequestration and orchestrating recovery, rather than simply tagging damaged substrates.

The identification of CHIP as a nucleolar organiser builds on its well-established role as an HSP70 co-chaperone and Ub ligase. At physiological levels, CHIP acts with HSP70 to facilitate nucleolar storage capacity, whereas overexpression exaggerates this storage mode and delays recovery, consistent with its known ability to attenuate HSP70 refolding activity^13,55^. This dual role supports the idea of CHIP as a rheostat that biases HSP70 from refolding towards storage and ubiquitination, thereby stabilising stress-engaged states until clearance becomes possible. VCP, by contrast, provides the driving force for extraction of ubiquitinated proteins and governs their flux between compartments^56^. Inhibition of VCP impaired nucleolar ubiquitination and blocked recovery, extending its known roles in ERAD and chromatin clearance to nucleolar PQC^57^. Recent work on localised Ub signalling in non-cytosolic compartments^58^ has emphasised the importance of VCP linkage specificity, particularly in processing K63-linked chains. Our findings are consistent with this and suggest that VCP is not merely downstream of CHIP but functions as a central switch that both controls recovery of ribosome biogenesis and remodels CHIP-Ub foci, highlighting reciprocal regulation between the two.

The nucleolus adapts to stress through distinct morphological states that correspond to functional priorities. The free-flow morphology reflects a dynamic liquid phase supporting rRNA synthesis and rapid molecular exchange. The peripheral state appears to represent a partial reorganisation, where the nucleolar core becomes a PQC-enriched storage zone with altered viscosity, while biosynthetic machinery is displaced to the rim, permitting residual rRNA flux. By contrast, the sealed morphology indicates a pathological phase transition towards a gel-like state that entraps components, severely restricting mobility and delaying recovery. This framework is consistent with the modern view of the nucleolus as a biomolecular condensate formed by liquid-liquid phase separation, where material properties such as viscosity and surface tension dictate function^59,60^. Similar to stress granules, which can undergo aberrant solidification when overloaded with misfolded proteins^61^, sealed nucleoli likely represent a protective but maladaptive state when extraction mechanisms are saturated. Comparable structural-functional coupling has been described in *C. elegans*, where nucleolar stress bodies (NoSBs) arise upon inhibition of rRNA transcription or defects in processing^12^. These reorganisations, including ring-shaped nucleoli and vacuolisation, correlate with precursor accumulation and activation of global stress pathways. Our identification of free-flow, peripheral, and sealed morphologies mirrors this paradigm but extends it by linking structural transitions not only to transcriptional arrest but also to PQC flux, ubiquitination dynamics, and recovery of ribosome biogenesis.

In parallel, subcellular Ub proteomics has demonstrated that stress-induced ubiquitination is not uniformly distributed but enriched in distinct compartments, where adaptors and segregases provide selectivity^58^. We show that a subset of nucleolus-localized proteins becomes ubiquitinated during heat shock, and that both the timing of these modifications and their reversal by deubiquitination are critical for recovery. This supports a model in which nucleolar ubiquitination functions not merely as damage marking but as a structured signal embedded in a cycle of sequestration, retention, and release, coordinated by CHIP, VCP, and DUBs.

Finally, the influence of intrinsic nucleolar proteins such as NOL6 and WDR55 underscores the intimate coupling between biogenesis and proteostasis. NOL6 depletion reduced storage and accelerated recovery, suggesting that it normally provides a scaffold that stabilises rRNA metabolism and prolongs sequestration. WDR55 depletion produced the opposite phenotype, enforcing hyper-sequestration and delaying recovery. These divergent outcomes indicate that ribosome biogenesis machinery does not simply suffer collateral disruption during stress but actively feeds back into proteostasis by tuning the material state of the nucleolus. Even partial depletion of WDR55 was sufficient to alter nucleolar behaviour, consistent with its classification as a common essential gene in cancer cells^62^. Together, these results reinforce that nucleolar PQC, and ribosome biogenesis are inseparable processes, co-regulated through structural and functional interdependencies between nucleolar proteins and stress-responsive factors.

The integration of PQC with biosynthesis involves trade-offs that reflect the competing priorities of clearance and growth. Cells must decide whether to allocate resources to removing damaged proteins or to resuming rRNA transcription and ribosome production. CHIP overexpression or HSP70 inhibition biased nucleoli towards storage at the expense of recovery, while VCP inhibition or HSP70 activation allowed premature rRNA synthesis despite incomplete clearance. This suggests that the system is wired to balance risk: early resumption of rRNA synthesis may restore biosynthetic output but at the cost of sustaining a proteotoxic burden. ISR provides a critical safeguard in this balancing act. By globally repressing translation through eIF2α phosphorylation^63^, ISR reduces the influx of new misfolded clients, giving the nucleolus time to resolve stored substrates. Blocking ISR with ISRIB trapped CHIP in nucleoli during recovery, underscoring that nucleolar clearance is contingent on coordination with translational checkpoints.

The interplay with other stress pathways highlights that nucleolar PQC is not an isolated mechanism but part of an integrated proteostasis network. Tunicamycin-induced ER stress diverted CHIP away from nucleoli, illustrating triage of PQC resources toward the most acute crisis. Perturbations in nucleolar ubiquitination triggered rerouting of Bystin to cytoplasmic stress granules, linking nuclear and cytoplasmic compartments in a coordinated response. Crosstalk with long non-coding RNAs, particularly IGS lncRNAs that scaffold nucleolar detention centres^64^, offers an additional axis of regulation: these RNAs may act as selective retention platforms, modulating accessibility of PQC clients to VCP and determining the reversibility of storage.

Several limitations of our study should be considered. While we observe robust CHIP- and Ub-positive foci that correlate with delayed recovery, we cannot unambiguously distinguish misfolded clients such as DRiPs from stalled ribosome biogenesis complexes as the major retained substrates. Our analyses of VCP function relied on localisation and inhibitor-based perturbations rather than direct measurements of its ATPase or segregase activity and thus may not fully capture how VCP engages substrates within dense or sealed condensates. Although NOL6 and WDR55 depletion produced strikingly opposite phenotypes, indirect contributions via altered rRNA metabolism cannot be excluded. Likewise, our morphological classifications reveal strong functional correlations; however, causal testing will require direct manipulation of nucleolar material properties. While CHIP catalytic activity clearly influenced nucleolar retention and recovery, we did not identify its substrates or the Ub chain types involved; the role of deubiquitinates as potential off-switches also remains to be validated. Finally, although we provide evidence of coupling between nucleolar PQC and ISR, ER stress, and stress granules, these links are inferred from perturbation outcomes and require mechanistic dissection.

In conclusion, our study establishes that nucleolar PQC functions as a continuum balancing folding, ubiquitination, sequestration, and ribosome biogenesis to sustain cellular fitness under stress. Disruption of this balance through defects in CHIP, VCP, or ribosome biogenesis factors may contribute to ribosomopathies, neurodegeneration, and ageing. These insights also suggest translational opportunities: targeting CHIP, VCP, or chaperone pathways could restore proteostasis or selectively destabilise malignant cells with hyperactive nucleoli, which are uniquely dependent on enhanced ribosome production^65–67^ Finally, the rRNA morphologies we define-free-flow, peripheral, and sealed-may serve as biomarkers of nucleolar resilience. These features are particularly relevant in light of recent evidence that mutant Huntingtin disrupts nucleolar integrity and rRNA metabolism in vivo^68^, suggesting that nucleolar PQC may represent a broader defence pathway against disease-associated protein aggregation.

## RESOURCE AVAILABILITY

### Materials availability

All unique materials and reagents generated in this study are available from the Lead Contact upon reasonable request. Requests for further information or resources should be directed to and will be fulfilled by Małgorzata Piechota or Wojciech Pokrzywa.

### Data and code availability

The RNA-seq datasets generated in this study have been deposited in the Gene Expression Omnibus (GEO) under accession number GSE190142. This paper does not report original code. Any additional information required to reanalyse the data reported in this paper is available from the Lead Contact upon request.

## Supporting information

STAR Methods Table

## ACKNOWLEDGMENTS

We thank the Genome Engineering Unit of the International Institute of Molecular and Cell Biology in Warsaw (IIMCB) for generating DNA constructs, the Microscopy and Cytometry Facility for support with confocal imaging, and the Bioinformatics Facility for assistance with RNA-seq data analysis. We are grateful to Aleksandra Szybińska for optimising HSP70 knockdown in HeLa CHIP cells and to Anna Grabowska for optimising blotting procedures. Access to core facilities was supported by the IIMCB IN-MOL-CELL Infrastructure, funded by the European Union - NextGenerationEU under the National Recovery and Resilience Plan. IN-MOL-CELL also received support from the European Union through Horizon Europe (Project 101059801, RACE) and from the RACE-PRIME project implemented within the IRAP programme of the Foundation for Polish Science, co-financed by the European Union under the European Funds for Smart Economy 2021-2027 (FENG). We also thank members of the Pokrzywa laboratory for their helpful comments and feedback on the manuscript.

## Funding

This work was primarily supported by the National Science Centre, Poland, through the SONATA BIS grants no. 2021/42/E/NZ1/00190 to W.P. and 2018/30/E/NZ7/00614 to W.S. The early phases of the project were supported by the Foundation for Polish Science, co-financed by the European Union under the European Regional Development Fund (grant POIR.04.04.00-00-5EAB/18-00), and by the Norwegian Financial Mechanism 2014–2021, operated by the Polish National Science Centre under project contract no. 2019/34/H/NZ3/00691.

### Declaration of generative AI and AI-assisted technologies in the writing process

During the preparation of this work the authors used ChatGPT to improve the readability and language of the manuscript. After using this tool, the authors reviewed and edited the content as needed and take full responsibility for the content of the published article.

## AUTHOR CONTRIBUTIONS

**Małgorzata Piechota:** Conceptualization; Data curation; Formal analysis; Methodology; Investigation; Validation; Visualization; Supervision; Writing-original draft; Writing-review & editing. **Lilla Biriczová:** Formal analysis; Investigation; Methodology; Visualization; Konrad Kowalski: Formal analysis; Investigation; Methodology; Visualization; **Fernando Aprile-García:** Formal analysis; Methodology; Investigation, Visualization, Data curation **Marta Leśniczak-Staszak:** Formal Analysis, Methodology, Investigation, Visualization; **Natalia A. Szulc:** Formal analysis, Methodology; Writing-review & editing **Witold Szaflarski:** Data curation, Resources, Supervision **Ritwick Sawarkar:** Resources, Supervision **Wojciech Pokrzywa:** Conceptualization; Data curation; Formal analysis; Funding acquisition; Project administration; Resources; Supervision; Validation; Writing-original draft; Writing-review & editing.

## STAR⍰METHODs

## EXPERIMENTAL MODEL

Several immortalised human cell lines were used in this study. HeLa Flp-In T-REx, a modified cervical cancer-derived HeLa cell line and HEK293 Flp-In T-REx, a derivative of embryonic kidney HEK293 cells, were obtained as kind gifts from Dr. R. Szczesny. A stable HeLa EGFP-NPM1 line with tetracycline-inducible EGFP-NPM1 expression was generated using HeLa Flp-In T-REx cells. MCF7, a breast cancer line derived from pleural effusion of a patient with adenocarcinoma, was kindly provided by Prof. A. Żylicz. The HEK293T NLS-LG line, stably expressing GFP-tagged luciferase with a nuclear localisation signal, was a kind gift from Dr. M. S. Hipp. The HEK293T GFP-CBX2 line was obtained from Prof. V. van den Boom. Stable HeLa CHIP and HEK CHIP cell lines with tetracycline-inducible EGFP-CHIP expression were generated in our laboratory from HeLa Flp-In T-REx and HEK293 Flp-In T-REx, respectively. All cell lines were cultured in Dulbecco’s Modified Eagle’s Medium (DMEM; D6429, Sigma-Aldrich) supplemented with 10% heat-inactivated fetal bovine serum (FBS; F9665, Sigma-Aldrich) and 1% antibiotic–antimycotic (15240062, Thermo Fisher Scientific) at 37 °C in a humidified 5% CO_2_ incubator. For experiments requiring tight control of tetracycline-inducible expression, Tet-System Approved FBS (A4736201, Thermo Fisher Scientific) was used instead of standard FBS. To maintain selection, HeLa CHIP and HEK CHIP cells were cultured with blasticidin (10 µg/ml; ant-bl-05, Invivogen) and hygromycin B (50 µg/ml; 10687010, Thermo Fisher Scientific), while HEK293T NLS-LG cells were maintained with G418 (100 µg/ml; 10131035, Thermo Fisher Scientific). Expression of EGFP-CHIP in HeLa CHIP and HEK CHIP lines was induced with tetracycline (25 ng/ml) added at plating or one day prior to experiments. For heat shock experiments, cells were transferred to a humidified incubator set at 42 °C for 90 min unless otherwise indicated and returned to 37 °C for recovery. Cells were passaged using trypsin (T4049, Sigma-Aldrich). All cell lines were routinely tested for mycoplasma contamination using PCR-based assays.

## METHOD DETAILS

### Poly-L-lysine coating

HEK293T cells were seeded on glass coverslips or 8-well chambered slides (80826, Ibidi) precoated with poly-L-lysine solution (P4707, Sigma-Aldrich). Coating was performed for 1 h at 37 °C, followed by two washes with sterile PBS and drying under a laminar flow hood.

### Tetracycline preparation

Tetracycline (TET701, BioShop) was prepared following a published protocol^69^. Briefly, tetracycline was dissolved in 96% ethanol to a concentration of 5 mg/ml and rotated for 30 min at room temperature. The solution was then incubated overnight at −20 °C, followed by an additional 30 min rotation the next day. After filtration through a 0.22 μm syringe filter, the stock was diluted with ethanol to a final concentration of 100 μg/ml and stored at −20 °C.

### Plasmid construction

Vectors pKK-EGFP-TEV, pKK-mCherry-TEV, and pKK-FLAG-TEV were kindly provided by Dr. R. Szczesny, and CHIP/CHIP K30A sequence templates by Dr. M. Figiel. All plasmids (mCherry-CHIP, mCherry-CHIP H260Q, mCherry-CHIP K30A, EGFP-CHIP, EGFP-CHIP H260Q, pKK-FLAG-HSPA1A, and HaloTag-ANXA5) were generated using sequence- and ligation-independent cloning (SLIC). For cloning, parental vectors (pKK-EGFP-TEV, pKK-mCherry-TEV, and pKK-FLAG-TEV) were linearised with BshTI and NheI, while pHTN-HaloTag was linearised with EcoRI. Linearised vectors were combined with PCR-amplified human target sequences, treated with T4 DNA polymerase, and transformed into bacteria. Primer sequences used for amplification, site-directed mutagenesis, and validation are listed in Table S4.

### Stable cell line generation

HeLa Flp-In T-REx and HEK293 Flp-In T-REx cells were seeded in 6-well plates and co-transfected with 1 μg pOG44 (a gift from Dr. R. Szczesny) and 0.8 μg EGFP-CHIP plasmid. Transfections were performed using Mirus reagents: 2 μl TransIT-293 (MIR 2700, Mirus) for HEK293 Flp-In T-REx, and 2 μl TransIT-HeLa together with 1.3 μl Monster (MIR 2900, Mirus) for HeLa Flp-In T-REx cells. One day after transfection, cells were placed under selection with 10 μg/ml blasticidin and 50 μg/ml hygromycin B, which was maintained for ~4 weeks until stable clones were established.

### Cell transfection

Transient transfections were carried out using Lipofectamine 2000 (E2312, Thermo Fisher Scientific) according to the manufacturer’s instructions. Cells were seeded on 8-well chambered slides (Ibidi) at a density of 5,000–20,000 cells/well and transfected with 200–500 ng plasmid DNA. A constant Lipofectamine:DNA ratio of 2:1 (μl:μg) was maintained. Transfections were performed ~24 h before imaging.

### HSP70 knockdown

HeLa CHIP cells were seeded on 35 mm imaging dishes (Ibidi) at a density of 200,000 cells. The following day cells were co-transfected with 75 pmols HSP70 siRNAs (IDs 145248 and 6965, Thermo Fisher Scientific) using 9 μl Lipofectamine RNAiMAX reagent (13778150, Thermo Fisher Scientific) in Opti-MEM Reduced Serum Medium (11058021, Thermo Fisher Scientific). The medium was exchanged after 48 h post-transfection. Silencing lasted 72 h.

### CHIP knockdown

For RNA sequencing, 1.5 × 10^5^ HeLa Flp cells were seeded per well in a 6-well plate. Transfection was performed 24 h after seeding using 7 μl Lipofectamine RNAiMAX (13778150, Thermo Fisher Scientific) and 50 pmol CHIP siRNA (ID 195025, Thermo Fisher Scientific) or negative control siRNA #2. siRNAs were diluted in 125 μl Opti-MEM (11058021, Thermo Fisher Scientific) and combined with Lipofectamine diluted in 125 μl Opti-MEM. After a 5-min incubation at room temperature, 250 μl siRNA-lipid complexes were added per well. Cells were harvested after 48 h of silencing.

For nucleolar morphology analysis, 3 × 10^4^ HeLa Flp cells were seeded on coverslips in 24-well plates. Cells were co-transfected with 10 pmol CHIP siRNA (or negative control #2) using 1.4 μl Lipofectamine RNAiMAX in 50 μl Opti-MEM. Cells were analysed 48 h post-transfection.

### RNAi miniscreen

HeLa Flp cells were seeded at a density of 1 × 10^4^ cells per well in 8-well chambered slides (80826, Ibidi). Forward transfection was performed with 3 pmol siRNA (target sequences listed in Table S5) and 0.7 μl Lipofectamine RNAiMAX (13778150, Thermo Fisher Scientific) in Opti-MEM Reduced Serum Medium (11058021, Thermo Fisher Scientific). Cells were maintained for 48 h of silencing, after which they were subjected to the heat shock assay. This transfection formulation was subsequently used for targeted depletion of WDR55 and NOL6 in follow-up experiments.

### Compound treatment

All compounds, except Leptomycin B (LMB, dissolved in methanol:water, 7:3), were prepared as stock solutions in DMSO. For heat shock assays, cells were typically pretreated with compounds before 90 min heat shock, unless otherwise indicated, and treatments were maintained throughout the recovery period. Details of compound identity, concentrations, and exposure times are provided in Table S6.

### Immunofluorescence

Cells were fixed in 4% formaldehyde for 10 min or 3.7% formaldehyde for 15 min at room temperature. Fixatives were prepared in PBS from 16% w/v formaldehyde ampules (28906, Thermo Fisher Scientific). Following fixation, cells were washed three times in PBS and permeabilised with 0.1% (v/v) Triton X-100 in PBS for 10 min at room temperature. Samples were incubated in blocking buffer (2% BSA, 1.5% goat serum, 0.1% Triton X-100 in PBS) for 10-30 min at room temperature. Primary antibodies (Table S7) were diluted in blocking buffer and incubated overnight at 4 °C. After PBS washes, appropriate fluorescent secondary antibodies (1:500 dilution, Table S7) were applied for 60 min at room temperature. Cells were washed three times in PBS (5 min each). Coverslip-grown samples were mounted in Vectashield antifade mounting medium with DAPI (H-1800-10, Vector Laboratories). For nuclei visualisation with custom polyvinyl mounting medium, cells were incubated with 4 μM Hoechst 33342 (62249, Thermo Fisher Scientific) for 10 min during the second PBS wash. For cells cultured in 8-well chambered slides, PBS was retained after the final wash and samples were directly imaged.

### FISH (Fluorescence in situ hybridization)

Approximately 50,000 cells were grown on coverslips, fixed with 4% paraformaldehyde in PBS (sc-281692, Santa Cruz Biotechnology) for 10 min, and permeabilised in cold 96% methanol for 10 min. FISH was performed essentially as described^70^. Briefly, samples were blocked in PerfectHyb Plus Hybridisation Buffer (H7033, Sigma-Aldrich) for 15 min at 52 °C and hybridized with a Cy5-labelled 47S rRNA probe for 1 h at 52 °C. After hybridisation, samples were washed twice in 2× SSC (15557044, Thermo Fisher Scientific) and once in PBS. Cells were subsequently incubated with primary and secondary antibodies (Table S7) and DAPI (D9542, Sigma-Aldrich), each for 45 min. Finally, coverslips were washed twice in PBS and mounted in polyvinyl mounting medium. Images were acquired on an Olympus FV10i laser scanning confocal microscope and analysed parametrically using Imaris software.

### HaloTag-RPL11 and HaloTag-ANXA5 labeling

Labelling was performed essentially as described previously^71^. Cells grown in 8-well chambered slides were transfected with 200 ng HaloTag-ANXA5 or HaloTag-RPL11 plasmids, and heat shock assays were carried out the following day. For general labelling of fusion proteins, cells were incubated with 0.7 µl Janelia 646 HaloTag ligand (JF646, GA1121, Promega; 0.6 µM final concentration) for 15 min before heat shock, followed by two PBS washes. To selectively label nascent HaloTag-RPL11, pre-existing HaloTag was first blocked with 100 µM 7-Bromo-1-heptanol (7BRO, H54762.03, VWR) for 15 min, followed by three PBS washes (2 min each with shaking). Fresh medium was then added containing 0.5 µl JF646 (0.4 µM final) and the indicated compounds. After the heat shock assay, cells were washed twice in PBS (2 min each) prior to fixation.

### OPP labeling combined with mCherry-CHIP transfection

HeLa Flp cells seeded on coverslips in 24-well plates were transiently transfected with mCherry-CHIP plasmids, and expression was induced with 25 ng/ml tetracycline. The following day, cells were treated with 25 µM O-propargyl-puromycin (OPP, 10456, Thermo Fisher Scientific) and subjected to heat shock for 2 h. For recovery, heat-shocked cells were returned to 37 °C for 1–2 h, with OPP treatment maintained throughout. OPP labelling was carried out according to the manufacturer’s protocol. Briefly, cells were fixed in 3.7% formaldehyde (15 min) and permeabilised in 0.5% Triton X-100 (15 min, room temperature). After two PBS washes, incorporated OPP was conjugated with Alexa Fluor 488 azide by click chemistry. Samples were subsequently washed in rinse buffer and PBS and mounted for imaging.

### OPP labeling combined with NOL6 and WDR55 silencing

HeLa Flp cells (10,000 per well) were seeded in 8-well chambered slides. The following day, cells were transfected with 3 pmol siRNAs targeting NOL6 or WDR55 using Lipofectamine RNAiMAX in 10 µl Opti-MEM. Control cells were transfected with an equivalent amount of scrambled siRNA. Cells were cultured for an additional 48 h prior to the heat shock assay and O-propargyl-puromycin (OPP) labelling (10457, Thermo Fisher Scientific). Cells were subjected to heat shock for 90 min, then treated with 20 µM OPP. One slide remained under heat stress for an additional 1 h, while the other was transferred to 37 °C for 1 h recovery. Control cells (no heat stress) were incubated with 20 µM OPP for 1 h at 37 °C. Following treatment, cells were washed twice in PBS, fixed with 3.7% formaldehyde (15 min, room temperature), and permeabilised with 0.5% Triton X-100 (15 min, room temperature). After two PBS washes, incorporated OPP was conjugated with Alexa Fluor 594 azide by click chemistry, according to the manufacturer’s protocol. Samples were then washed in rinse buffer and PBS before proceeding with immunofluorescence for mono- and polyubiquitinated conjugates.

### 5-ethynyl uridine (EU) labelling

EU labelling was performed analogously to OPP labelling (10330, Thermo Fisher Scientific). 10,000 HeLa CHIP cells were seeded in 8-well chambered slides. The following day, cells were transfected with 3 pmol siRNAs targeting NOL6 or WDR55 using Lipofectamine RNAiMAX in 10 µl Opti-MEM. Control cells were transfected with an equivalent amount of scrambled siRNA. For CHIP induction, cells were treated with 25 ng/ml tetracycline one day before the assay. In experiments testing compound effects, cells were instead pretreated with tunicamycin, VER, CB-5083, or SW02, which were maintained during heat shock and recovery. Cells were subjected to 90 min heat shock at 42 °C, after which they were incubated with 1 mM EU (5-ethynyl uridine) either (i) under heat shock in fresh pre-heated medium or (ii) at 37 °C for 1 h recovery. Control cells not subjected to heat stress were incubated with 1 mM EU for 1 h at 37 °C. Following treatment, cells were washed twice in PBS, fixed with 3.7% formaldehyde (15 min, room temperature), and permeabilised with 0.5% Triton X-100 (15 min, room temperature). After two PBS washes, incorporated EU was conjugated with Alexa Fluor 594 azide by click chemistry according to the manufacturer’s protocol (30 min labelling). Samples were then washed in rinse buffer and PBS, followed by NPM1 immunofluorescence.

### Labelling with the proteasome activity sensor Me4BodipyFL-Ahx_3_Leu_3_VS

HeLa Flp cells (10,000 per well) were seeded into 8-well chambered slides. The next day, cells were transfected with 3 pmol siRNAs targeting NOL6 or WDR55 using Lipofectamine RNAiMAX in Opti-MEM. Control cells received an equivalent amount of scrambled siRNA. Following transfection, cells were incubated for 48 h before the heat shock assay. Cells were subjected to 90 min heat shock at 42 °C, after which the medium was replaced with fresh preheated medium (42 °C) containing 1 μM Me4BodipyFL-Ahx_3_Leu_3_VS proteasome activity sensor (Me4) (prepared as 1 μl sensor stock per 250 μl medium; I-190, Bio-Techne). For recovery experiments, cells were exposed to 1 μM Me4 sensor and transferred immediately to 37 °C for 1 h. Control cells not subjected to heat stress were incubated with 1 μM Me4 sensor for 1 h at 37 °C. After treatments, cells were washed twice with PBS, fixed in 3.7% formaldehyde (15 min, room temperature), and processed for NPM1 immunofluorescence.

### Leptomycin B-based Bystin retention assay

Leptomycin B (LMB) treatment and interpretation of Bystin localisation followed published guidelines with minor modifications^53^. HeLa Flp cells (10,000 per well) were seeded into 8-well chambered slides and transfected with 3 pmol siRNAs targeting NOL6 or WDR55 using Lipofectamine RNAiMAX. Control cells were transfected with an equivalent amount of scrambled siRNA. For heat shock experiments, cells were exposed to 42 °C for 2 h in the presence of 10 nM LMB. For recovery experiments, cells were first subjected to 2 h heat shock, followed by replacement with fresh medium containing 10 nM LMB and immediate transfer to 37 °C for 1 h recovery. Control cells not subjected to heat stress were incubated with 10 nM LMB at 37 °C for 2 h. After treatments, cells were fixed and immunostained for Bystin.

### Image acquisition

Confocal microscopy was performed using a ZEISS LSM800 laser scanning confocal microscope (Carl Zeiss Microscopy) equipped with 63×/1.4 NA and 40×/1.3 NA oil immersion objectives. Unless otherwise indicated, images represent single optical sections. For colocalisation studies and enhanced resolution, imaging was performed with the Airyscan detector module. Image acquisition parameters (laser power, detector gain, and pixel size) were kept constant within each experiment to allow quantitative comparisons.

### Fluorescence recovery after photobleaching (FRAP)

Cells for FRAP experiments were cultured on 35-mm glass-bottom imaging dishes (Ibidi). FRAP measurements were performed on a ZEISS LSM800 confocal laser scanning microscope equipped with a 40×/1.3 NA oil-immersion objective. A circular region of interest (ROI) of constant size was selected within nucleoli, and bleaching was performed using 100% laser power of the 488-nm laser line. Fluorescence recovery was recorded for up to 3 min, with a frame interval of 0.5 s for CHIP or 1 s for NPM1. FRAP movies were analysed using FRAPAnalyser software (https://github.com/ssgpers/FRAPAnalyser). Fluorescence intensities were corrected for background signal and acquisition-related photobleaching.

### Nucleoli isolation

Nucleoli were isolated according to a previously described protocol with modifications^72^. Cells were cultured on 100-mm tissue culture dishes and harvested on ice. The medium was aspirated, and cells were washed once with 4 ml ice-cold PBS. Cells were then pelleted at 220 × g for 5 min at 4°C, resuspended in 5 ml Buffer A (10 mM HEPES pH 7.9, 10 mM KCl, 1.5 mM MgCl_2_, 0.5 mM DTT, 1× Complete protease inhibitor cocktail [04 693 159 001, Roche]), and incubated on ice for 5 min. Cells were homogenized on ice using a pre-chilled 1-ml Dounce homogenizer (Wheaton) with a tight pestle (10 strokes). The homogenate was centrifuged at 220 × g for 5 min at 4°C. The supernatant was collected as the cytoplasmic fraction. The pellet was resuspended in 3 ml S1 buffer (0.25 M sucrose, 10 mM MgCl_2_, 1× protease inhibitor cocktail), layered onto 3 ml S2 buffer (0.35 M sucrose, 0.5 mM MgCl_2_, 1× protease inhibitor cocktail), and centrifuged at 1430 × g for 5 min at 4°C. The pellet was resuspended in 3 ml S2 buffer to yield a nuclear suspension. To disrupt nuclei, suspensions were sonicated on ice (Sonica VCX130, ¼-inch tip) at 50% amplitude for 11 cycles (10 s pulse, 10 s rest). The sonicated sample was layered onto 3 ml S3 buffer (0.88 M sucrose, 0.5 mM MgCl_2_, 1× protease inhibitor cocktail) and centrifuged at 3000 × g for 10 min at 4°C. The supernatant was collected as the nucleoplasmic fraction. The pellet was resuspended in 500 µl S2 buffer and centrifuged at 1430 × g for 5 min at 4°C. The final pellet containing nucleoli was resuspended in 500 µl S2 buffer and stored at −80°C until use.

### Estimation of the protein concentration and Western blotting

Protein concentration was determined using the Pierce Rapid Gold BCA Protein Assay Kit (A53225, Thermo Fisher Scientific) according to the manufacturer’s instructions. For immunoblotting, protein samples were prepared in reducing SDS loading buffer and resolved on 10% acrylamide gels using Tris-glycine running buffer (25 mM Tris, 190 mM glycine, 0.1% SDS). Electrophoresis was performed at 90 V for stacking and 150 V for separating gels. Proteins were transferred to PVDF membranes using wet transfer at 400 mA constant current for 1 h at 4°C in transfer buffer (25 mM Tris, 190 mM glycine, 20% methanol, pH 8.3). For visualizing proteins on a membrane No-Stain Protein Labeling Reagent (A44449, Thermo Fisher Scientific) was used according to the manufacturer’s guidelines. Membranes were blocked in 5% (w/v) skimmed milk in TBST (50 mM Tris, 150 mM NaCl, 0.1% Tween-20, pH 7.6) for 1 h at room temperature, followed by overnight incubation at 4°C with primary antibodies diluted in the same blocking solution. After three washes in TBST (10 min each), membranes were incubated with HRP-conjugated secondary antibodies (1:10,000) for 1 h at room temperature. Protein detection was performed using SuperSignal West Pico PLUS Chemiluminescent Substrate (34580, Thermo Fisher Scientific), and images were acquired with a ChemiDoc Imaging System (Bio-Rad).

### RNA sequencing

Total RNA was isolated using TRI Reagent (93289, Sigma-Aldrich) according to the manufacturer’s instructions, followed by quantification and quality assessment with a NanoDrop spectrophotometer (Thermo Fisher Scientific) and Bioanalyzer RNA 6000 Nano Kit (Agilent). Only RNA samples with an RNA integrity number (RIN) > 8.0 were used for library preparation. Libraries were prepared using the TruSeq Stranded mRNA Library Prep Kit (Illumina), which includes poly(A) selection and strand-specific library construction. Sequencing was carried out on an Illumina HiSeq 2500 system to generate paired-end 2 × 100 bp reads.

### Image analysis

Image analysis was performed primarily with ImageJ software (https://imagej.nih.gov/ij/index.html)^73^. In HeLa CHIP cells nucleoli were manually delineated, and CHIP (EGFP) intensity was quantified as the mean gray value. For colocalisation studies, the JaCoP plugin^74^ was applied. Background subtraction was carried out using the rolling ball algorithm (radius = 150). Thresholds for the green and red channels were defined manually and maintained consistently across all images within a given experiment. For selected datasets (Figures 4Dy, 6A2, 7A, B1, E2, F2, S4A2, S6B2), image analysis was performed using customised CellProfiler pipelines^75,76^ to ensure reproducibility and high-throughput quantification. During manuscript preparation, brightness and contrast were adjusted uniformly across entire images only for visualisation purposes, without altering quantitative data or interpretation.

### Imaris 3D reconstruction and parametric analysis of FISH images

The crude Z-stack FISH images were collected using an Olympus FV10i confocal laser scanning microscope. Then, images were processed with Imaris 7.4.2 (Bitplane, UK) for 3D reconstruction. The volume of the nucleus (DAPI), nucleoli (NPM1) and 47S-rRNA were automatically calculated based on Z-stacks composed of individual images.

### Differential gene expression analysis

Differential gene expression analysis was performed with DESeq2 R package^77^ with default parameters. Genes with an adjusted p-value < 0.05 were considered significantly differentially expressed. MA plots and Venn diagrams were generated with custom R/Bioconductor scripts, making use of the ggplot2^78^ and VennDiagram packages^79^. For enrichment analysis, protein-coding genes were classified as nucleolar or non-nucleolar according to the annotation of nucleolus-localized proteins in the Human Protein Atlas, version 24.0^80^.

### Pathway enrichment analysis and gene-pathway network generation

Gene set enrichment analysis was performed using the R package clusterprofiler^81^. Over-representation testing was carried out on Gene Ontology (GO) Biological Processes gene sets, with p-values adjusted by the Bonferroni method. The *minGSSize* parameter was set to 10 to exclude very small categories. Enrichment results were visualised as dot plots, showing aggregated outcomes across multiple comparisons to highlight differentially affected biological processes. Dot plots were generated using the compareCluster function from clusterProfiler combined with custom R/Bioconductor scripts, employing ggplot2^78^. To explore higher-order relationships, gene-pathway interaction networks were constructed using custom scripts in the R/Bioconductor environment, with visualisation performed using the igraph^82^, ggraph^83^ and rrvgo^84^ packages.

## QUANTIFICATION AND STATISTICAL ANALYSIS

Data were analysed and visualised using GraphPad Prism software. Statistical significance was assessed using one-way or two-way ANOVA followed by appropriate post hoc multiple comparisons tests. For the 5-EU assay, a Kruskal-Wallis test followed by Dunn’s multiple comparisons test was applied. For comparisons between two independent groups, either a two-tailed unpaired t-test or a Mann–Whitney test was used, depending on data distribution. Significance levels are denoted in the figures as follows: p < 0.05 (*), p < 0.01 (**), p < 0.001 (***), p < 0.0001 (****); ns, not significant. Unless otherwise indicated, n refers to the number of nucleoli analysed, with details provided in the figure legends. Each experiment was performed independently on 2-4 biological replicates conducted on separate days. Violin plots represent the median, quartiles, and full data distribution, pooled across two independent experiments or shown as a representative dataset, as specified in figure legends.

## SUPPLEMENTAL INFORMATION

**Document S1**. Figures S1-S7 and Tables S4-S7 (related to STAR Methods).

**Table S1**. Differential gene expression upon CHIP depletion in basal conditions (related to Figure 4E).

**Table S2**. Differential gene expression under heat-shock conditions in CHIP-depleted versus scrambled siRNA-treated cells (related to Figure 4F).

**Table S3**. Overrepresentation analysis of Gene Ontology (GO) Biological Processes gene sets (related to Figure 4G).

**Video S1**. Time-lapse imaging of CHIP (green) and RPL11 (red) exchange in budded nucleoli (related to Figure 3E). Scale bar, 5 µm.

---

**Fig. S1.**
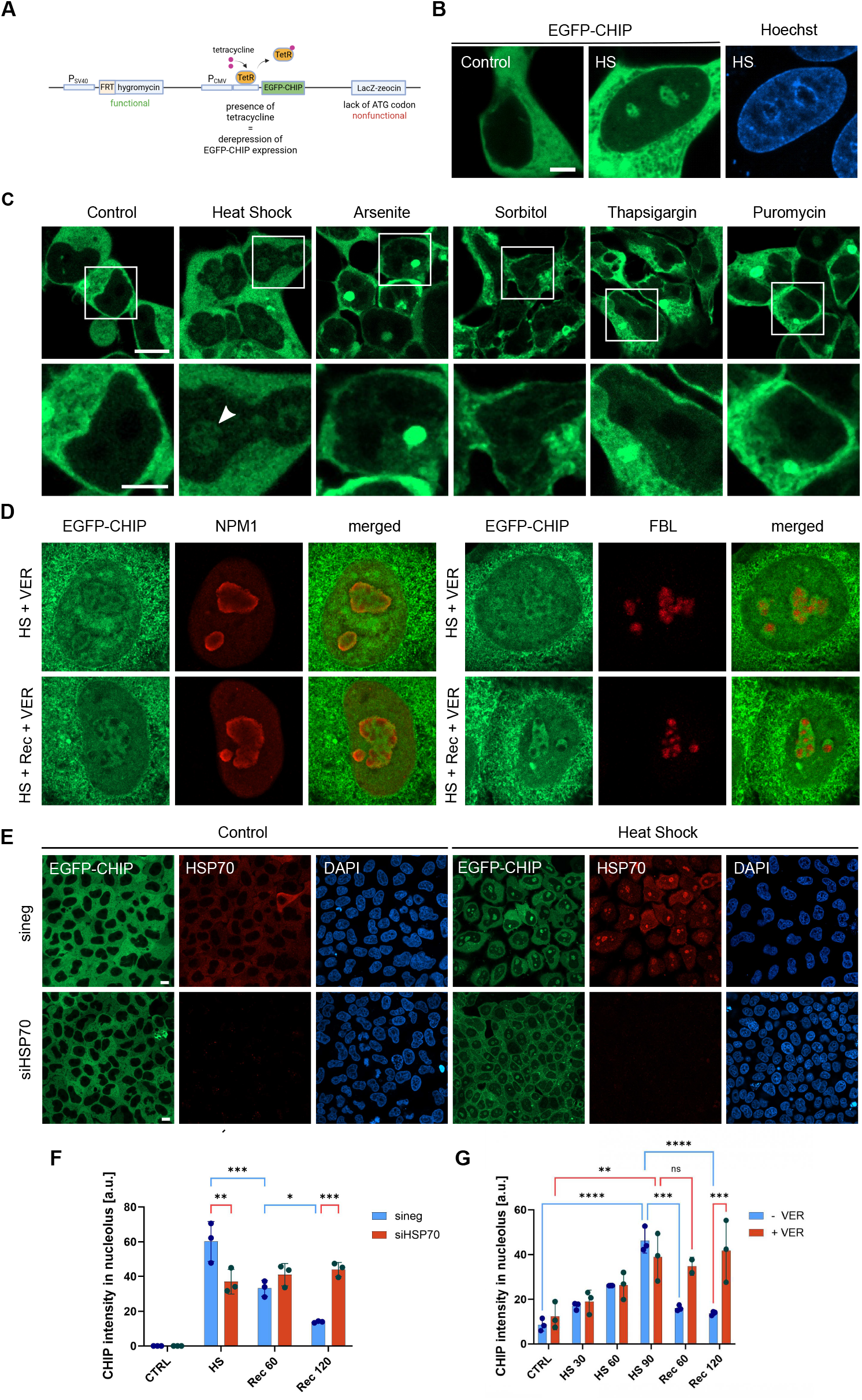

**Fig. S2.**
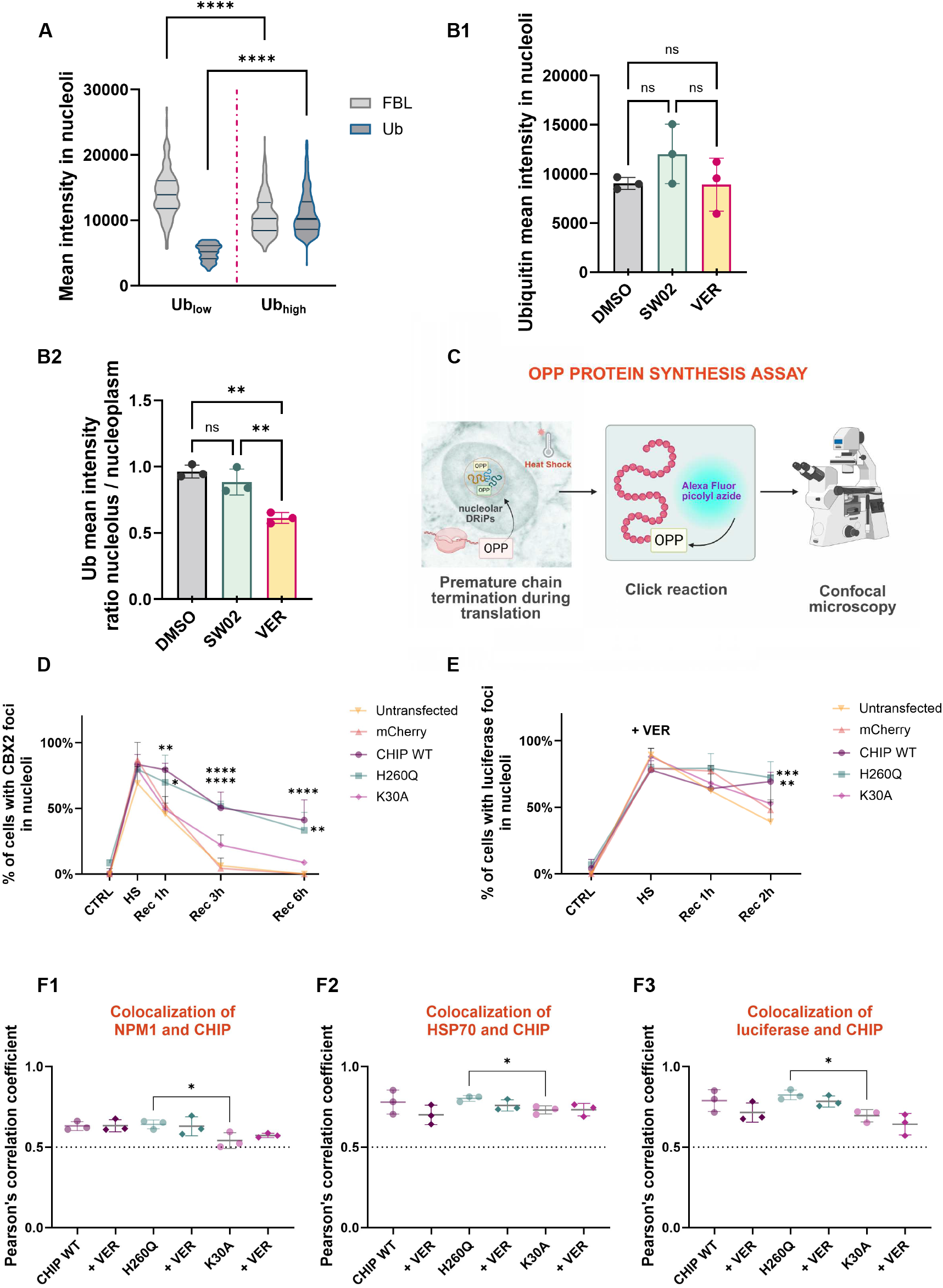

**Fig. S3.**
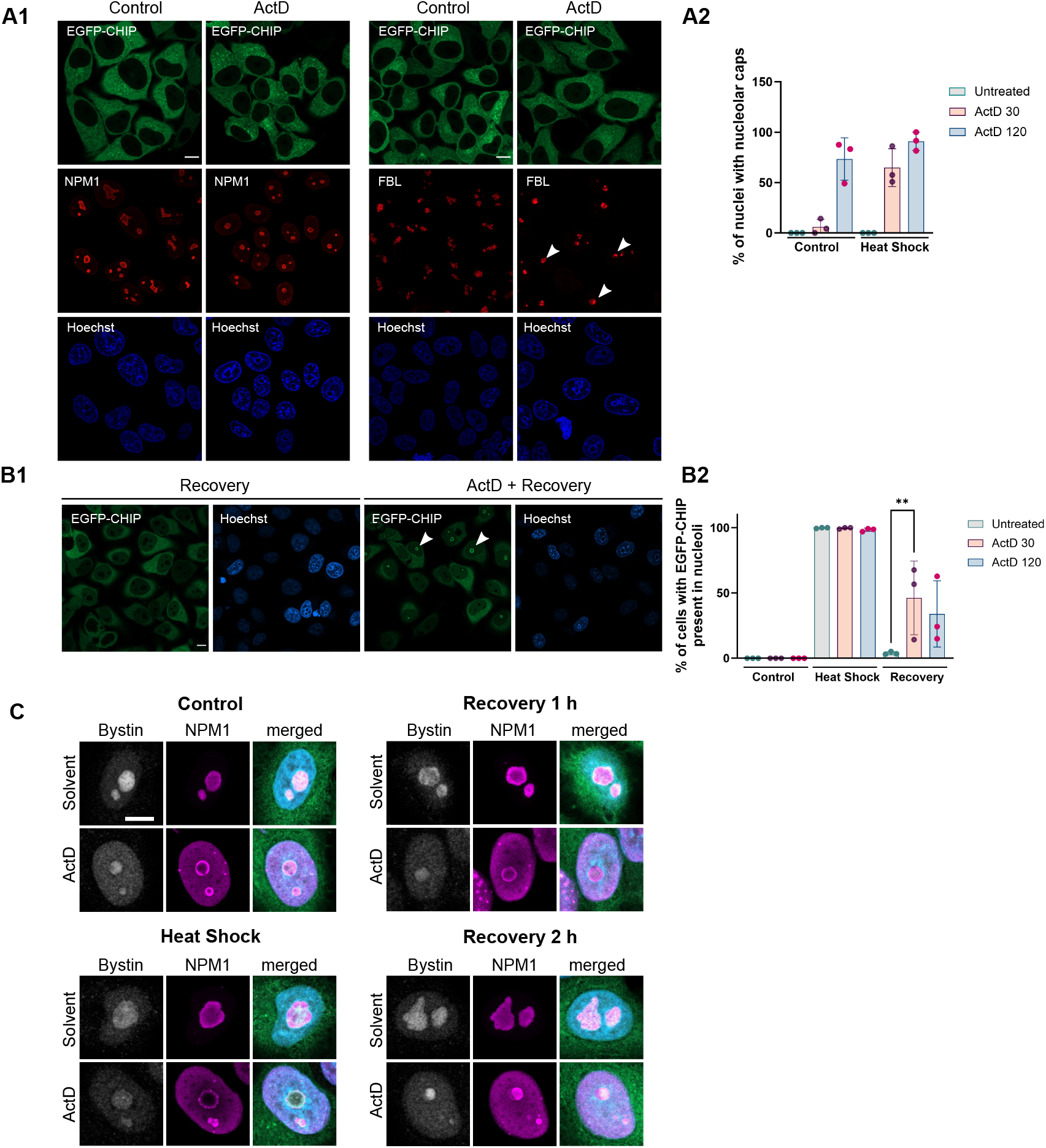

**Fig. S4.**
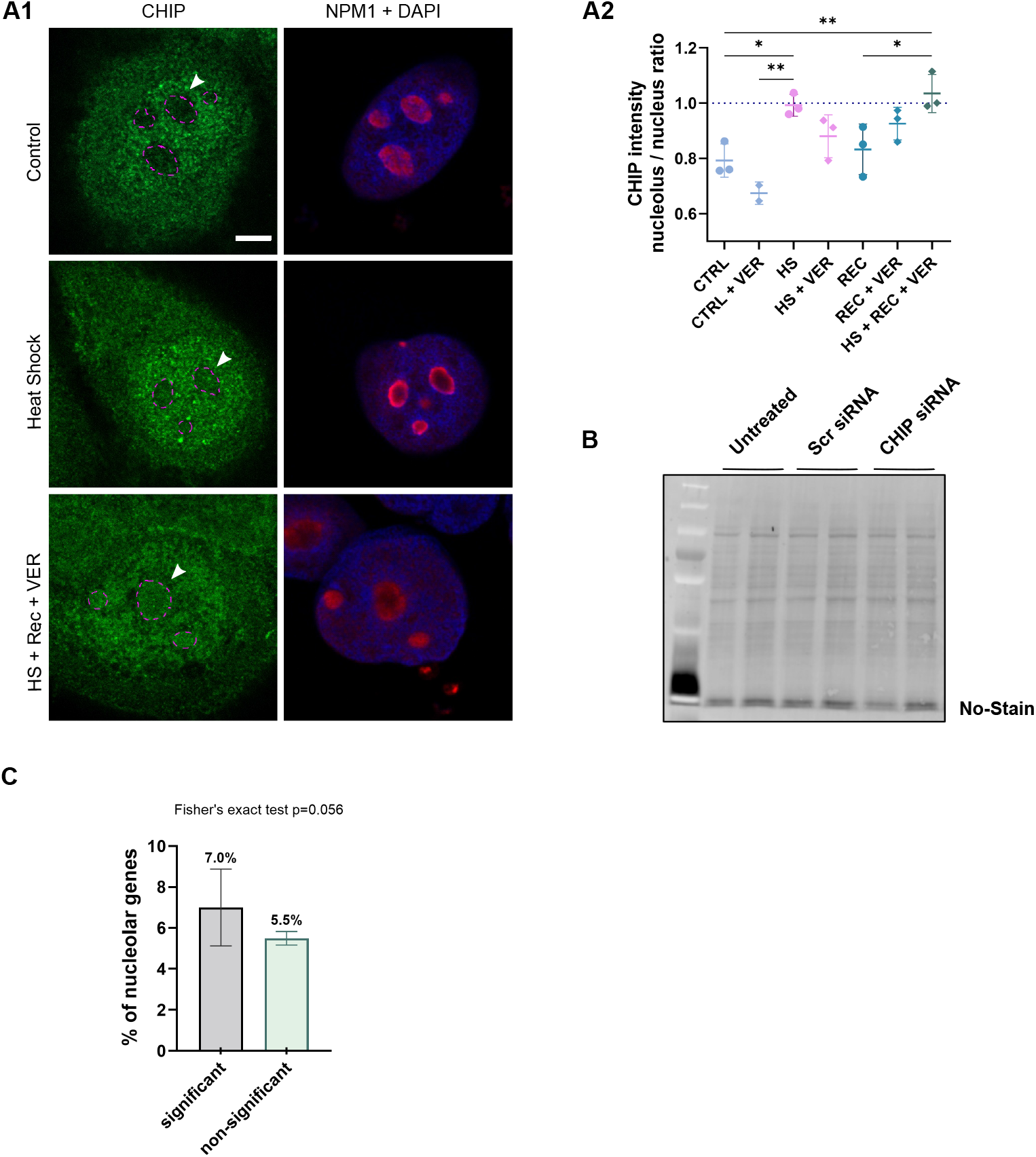

**Fig. S5.**
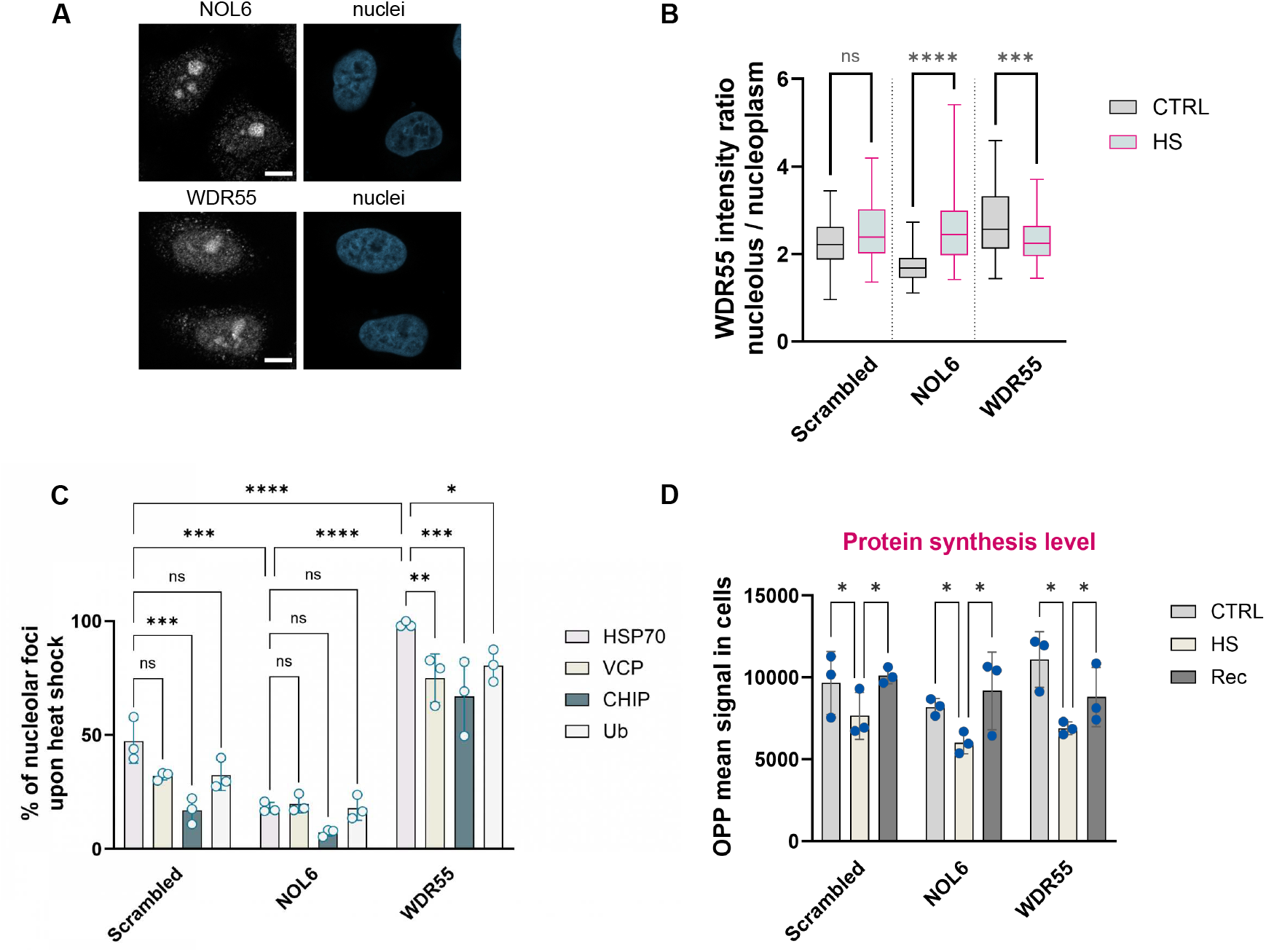

**Fig. S6.**
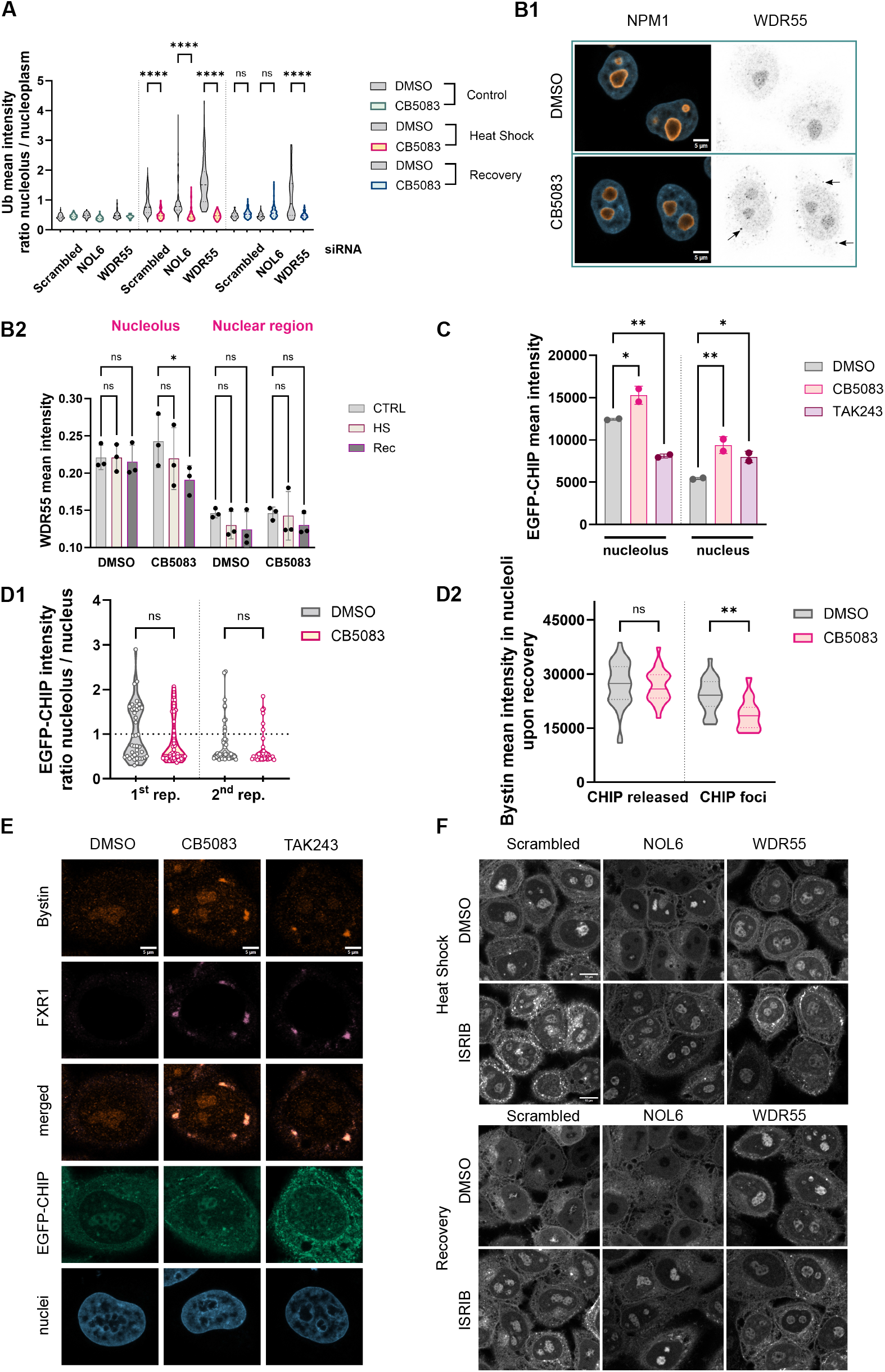

**Fig. S7.**
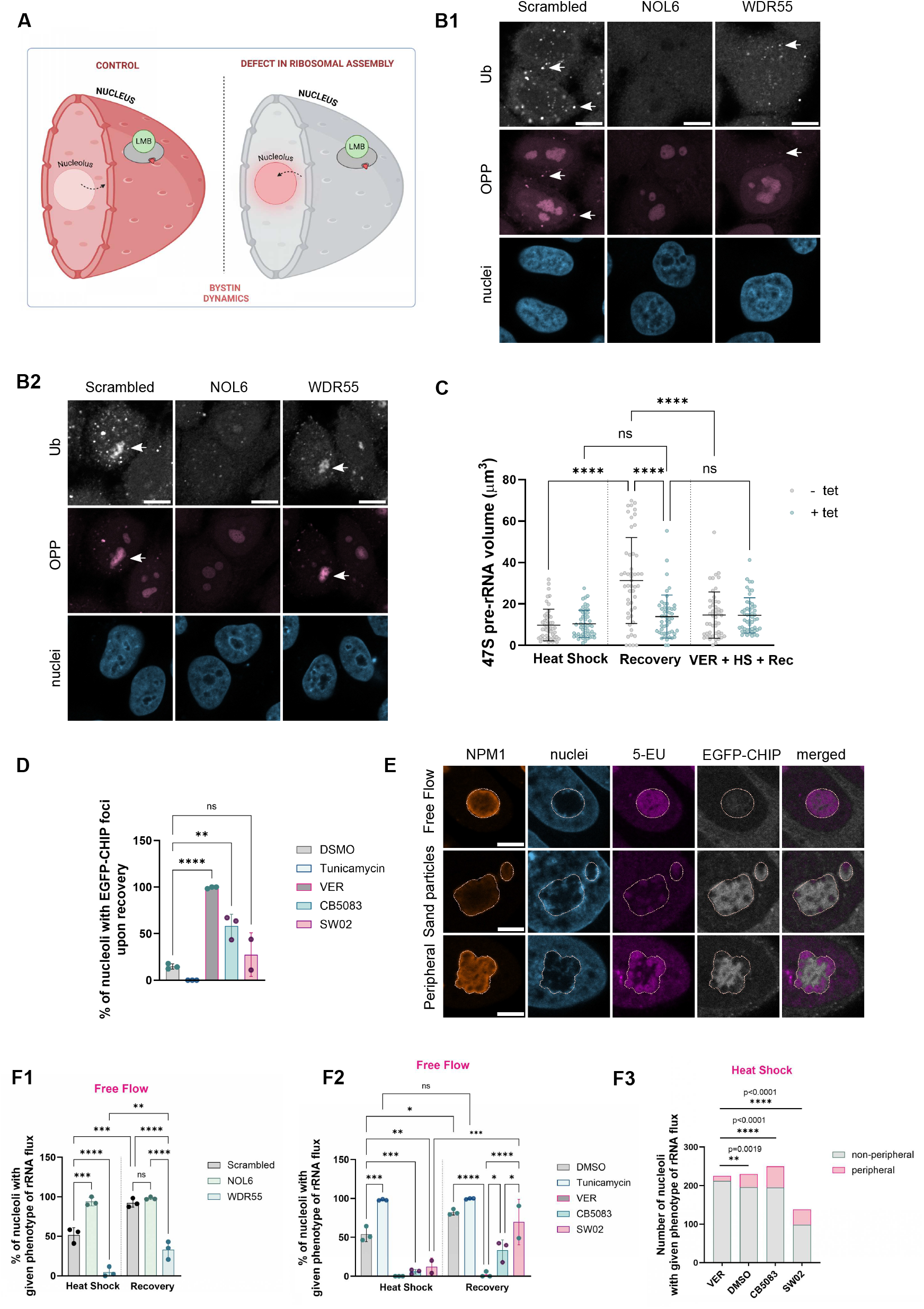

