## Supplemental Legends for "CHIP, VCP, and Nucleolar Gatekeepers Remodel the Nucleolus into a Stress-Responsive Proteostasis Hub"

**Figure S1. CHIP and HSP70 regulate nucleolar engagement and clearance during heat stress**

**(related to Figure 1).**

(A) Schematic of the EGFP-CHIP transgene integrated into HeLa Flp-In T-REx and 293 Flp-In T-REx cells (adapted from Szczesny et al., 2018). Cells are hygromycin B-resistant and zeocin-sensitive. EGFP-CHIP expression is TetR-repressed and induced by tetracycline.

(B) Confocal images of MCF7 cells transiently expressing EGFP-CHIP. Nucleoli identified by lack of Hoechst 33342 staining. Scale bar, 5 µm.

(C) CHIP nucleolar recruitment under heat shock conditions. HEK CHIP cells exposed to 42 °C (90 min), 50 µM sodium arsenite, 100 nM thapsigargin, 0.6 M sorbitol, or 2 mg/ml puromycin for 2 h. Arrowhead marks nucleolar CHIP in heat-shocked cells. Scale bars, 10 µm; magnified views, 5 µm.

(D) Confocal Airyscan images of HeLa CHIP cells immunostained for NPM1 (left) or FBL (right) after heat shock ± 40 µM VER. Cells treated with VER throughout heat shock and recovery are also shown. Scale bars, 5 µm.

(E) Confocal images of HeLa CHIP cells after HSP70 knockdown (siHSP70 vs non-targeting siRNA (sineg)). Cells were heat-shocked or left untreated and immunostained for HSP70. Nuclei stained with DAPI. Scale bar, 10 µm.

(F) Quantification of mean CHIP intensity in nucleoli during heat shock and recovery in control (sineg) and siHSP70 cells. Mean ± SD from n = 3 experiments. Two-way ANOVA with Tukey’s test.

(G) HSP70 inhibition by VER does not block CHIP entry into nucleoli during heat shock but delays release during recovery. CHIP intensity in nucleoli quantified after pretreatment with 40 µM VER. Mean ± SD from n = 3 experiments. Two-way ANOVA with Tukey’s test. Rec – recovery, the numbers next to it indicate the duration in minutes.

**Figure S2. CHIP expands nucleolar storage of misfolded proteins via HSP70 (related to Figure 2).**

(A) FBL levels decrease in Ub-positive nucleoli after heat shock. Violin plots show median and quartiles (n = 153 [Ub_low_], 291 [Ub_high_] nucleoli, 2 experiments). Welch’s t-test.

(B) VER alters nucleolar ubiquitination. (B1) Mean Ub intensity. (B2) Ub nucleolus/nucleoplasm ratio. Mean ± SD from n = 3 experiments. One-way ANOVA with Tukey’s test.

(C) Schematic of nucleolar DRiPs labelling by click chemistry.

(D) CHIP delays clearance of GFP-CBX2 foci. Percentage of nucleoli with GFP-CBX2 foci during heat shock and recovery ± CHIP WT, H260Q, or K30A. Mean ± SD from n = 3 experiments. Two-way ANOVA with Tukey’s test.

(E) VER inhibits luciferase clearance. Percentage of nucleoli with luciferase foci in the presence of VER and CHIP WT, H260Q, or K30A. Mean ± SD from n = 3 experiments. Two-way ANOVA with Tukey’s test.

(F) Pearson’s correlation between CHIP and NPM1 (F1), HSP70 (F2), or nucleolar luciferase (F3) during heat shock ± VER. Mean ± SD from n = 3 experiments. Welch’s t-test.

**Figure S3. Actinomycin D reveals coupling of nucleolar morphology, CHIP retention, and Bystin dynamics during stress recovery (related to Figure 3).**

(A) Actinomycin D (ActD) alters nucleolar morphology without inducing CHIP migration. (A1) Confocal images of HeLa CHIP cells after 2 h ActD treatment. Arrowheads mark FBL-positive nucleolar caps. Scale bar, 10 µm. (A2) Percentage of nuclei with nucleolar caps in basal and heat-shock conditions. Mean ± SD from n = 3 experiments.

(B) ActD pretreatment impairs CHIP release from nucleoli during recovery. (B1) Confocal images of HeLa CHIP cells after heat shock and recovery ± ActD. Arrowheads indicate nucleolar CHIP in ActD-treated cells. Scale bar, 10 µm. (B2) Percentage of cells with nucleolar CHIP after recovery, with ActD pretreatment for 30 min or 2 h. Mean ± SD from n = 3 experiments (one-way ANOVA with Dunnett’s test).

(C) ActD and heat shock co-treatment alters Bystin dynamics. Confocal images of HeLa CHIP cells stained for NPM1 and Bystin after heat shock and recovery ± ActD. Scale bar, 5 µm.

**Figure S4. CHIP relocalisation in MCF7 cells and enrichment of nucleolar genes among deregulated transcripts (related to Figure 4).**

(A) Endogenous CHIP relocalises to nucleoli upon heat shock in MCF7 cells. (A1) Representative confocal images of MCF7 cells in basal conditions, after heat shock, and after recovery in the presence of VER. Cells were immunostained for CHIP (green) and NPM1 (red), and nuclei counterstained with DAPI (blue). Arrowheads and dashed circles indicate nucleoli. Scale bar, 5 µm. (A2) Quantification of nucleolar versus nuclear CHIP intensity ratios. Mean ± SD from n = 3 experiments. One-way ANOVA with Tukey’s test.

(B) Protein visualization on the membrane with the No-Stain Reagent.

(C) Proportion of nucleolar genes among deregulated (significant) versus non-deregulated (non-significant) genes upon CHIP depletion. Bar plot shows the percentage of nucleolar genes in each group, with 95% confidence intervals.

**Figure S5. Nucleolar factors NOL6 and WDR55 modulate CHIP-dependent storage capacity**

**(related to Figure 5).**

(A) Endogenous NOL6 and WDR55 localise to nucleoli in HeLa Flp cells. Confocal immunofluorescence. Scale bars, 10 µm.

(B) WDR55 depletion alters its nucleolar distribution during heat stress. Quantification of WDR55 nucleolar/nucleoplasmic intensity ratio under basal and heat-shock conditions after RNAi. Box-and-whisker plots (min to max). One-way ANOVA with Tukey’s test; n = 3 experiments.

(C) NOL6 knockdown reduces, while WDR55 depletion enhances, nucleolar recruitment of HSP70, VCP, CHIP, and ubiquitinated proteins under heat stress. Percentage of nucleolar foci (intensity ratio nucleolus/nucleoplasm > 1). Mean ± SD from n = 3 experiments. Two-way ANOVA with Tukey’s test.

(D) Global translation is unaffected by NOL6 or WDR55 knockdown. OPP incorporation and click-labelling of nascent peptides were quantified under basal, heat shock, and recovery conditions. Mean ± SD from n = 3 experiments. Two-way ANOVA with Tukey’s test.

**Figure S6. VCP and ISR Coordinate Nucleolar PQC with Cellular Proteostasis (related to Figure 6).**

(A) VCP inhibition disrupts nucleolar ubiquitination. Quantification of Ub nucleolar/nucleoplasmic ratios in control, heat shock, and recovery conditions ± CB5083. Violin plot shows median and quartiles (n = 357 [CTRL], 790 [HS], 541 [Rec] nucleoli). One-way ANOVA with Sidak’s test.

(B) VCP inhibition alters WDR55 localisation, redistributing it from nucleoli to cytoplasmic foci. (B1) Representative confocal images of heat-shocked cells ± CB5083. Cells immunostained for NPM1 (orange), WDR55 (grey, inverted LUT), and Hoechst 33342 (blue). Arrows mark cytoplasmic WDR55 foci. Scale bar, 5 µm. (B2) Quantification of WDR55 levels in nucleoli and nucleoplasm under basal, heat-shock, and recovery conditions ± CB5083. Mean ± SD from n = 3 experiments. Two-way ANOVA with Tukey’s test.

(C) CB5083 and TAK243 treatment alters EGFP-CHIP distribution between nucleoli and nucleoplasm upon heat shock. Mean ± SD from n = 2 experiments. Two-way ANOVA with Sidak’s test.

(D) VCP inhibition during recovery does not delay CHIP release but affects Bystin restoration. (D1) Quantification of EGFP-CHIP nucleolar/nuclear ratios during recovery ± CB5083. Violin plots show all data points (n = 34-49 nucleoli/condition; 2 experiments). Welch’s t-test. (D2) Quantification of Bystin levels in nucleoli with or without EGFP-CHIP. Violin plots show median and quartiles (n = 15-63 nucleoli/condition; 2 experiments). One-way ANOVA with Tukey’s test.

(E) Bystin redistributes to stress granules under CB5083 and TAK243 pretreatment. Confocal images showing Bystin colocalisation with FXR1 (stress granule marker) in HeLa CHIP cells after heat shock. Scale bars, 5 µm.

(F) ISR inhibition by ISRIB traps EGFP-CHIP in nucleoli of NOL6-depleted cells during heat shock and recovery. Confocal images of HeLa CHIP cells (EGFP-CHIP, grey) under NOL6 and WDR55 knockdowns ± ISRIB. Scale bars, 10 µm.

**Figure S7. Nucleolar Reorganisation Couples PQC Engagement with Ribosome Biogenesis (related to Figure 7).**

(A) Schematic of Bystin behaviour with leptomycin B (LMB): in control cells (left, normal 40S assembly) versus cells defective in 40S assembly (right).

(B) NOL6 knockdown reduces DRiP foci colocalising with Ub, both in basal and heat shock conditions. (B1) Confocal images under basal conditions. Arrows mark OPP- and Ub-enriched cytoplasmic foci. (B2) Confocal images under heat shock. Arrows mark OPP- and Ub-enriched nucleolar foci. Scale bars, 10 µm.

(C) CHIP overexpression and VER suppress rRNA derepression during recovery. Quantification of 47S pre-rRNA volumes by FISH. Mean ± SD (n = 50 nucleoli/condition). One-way ANOVA with Tukey’s test.

(D) HSP70 inhibition blocks CHIP clearance during recovery. Percentage of nucleoli retaining CHIP foci after recovery ± indicated compounds. Mean ± SD from 2-3 experiments. One-way ANOVA with Dunnett’s test.

(E) Distinct rRNA flux patterns correspond to nucleolar PQC states. Representative confocal images of nucleoli after heat shock showing free-flow, (sand particles) sealed or peripheral morphologies. Cells pulsed with 5-EU to label nascent rRNA. Dashed lines mark nucleoli. Scale bars, 5 µm.

(F) rRNA flux correlates with nucleolar recovery progression. (F1-F2) Percentage of free-flow phenotype after heat shock and recovery ± RNAi or compounds. Mean ± SD from 2-3 experiments. One-way ANOVA with Tukey’s test. (F3) Occurrence of the peripheral phenotype during heat shock compared between VER and other treatments. Fisher’s exact test.

**Table S4. List of primers used for cloning and sequencing.**

**Table S5. List of siRNAs used in this paper.**

**Table S6. List of compounds used in this paper with description of their application.**

**Table S7. List of antibodies used in this paper.**
