## Supplementary material for "CHIP, VCP, and Nucleolar Gatekeepers Remodel the Nucleolus into a Stress-Responsive Proteostasis Hub": Table S4

| **Target plasmid** | **Primer for amplification** | **Sequence** |
| --- | --- | --- |
| EGFP-CHIP WT  mCherry-CHIP WT  mCherry-CHIP K30A | hCHIP forward | GGATCCgaaaacctgtacttccaaggaACCGGTATGAAGGGCAAGGAGGAGAAG |
|  | hCHIP reverse | GATATCaccctgaaaatacaaattctcGCTAGCTCAGTAGTCCTCCACCCAGC |
| EGFP-CHIP H260Q  mCherry-CHIP H260Q | hCHIP forward | GGATCCgaaaacctgtacttccaaggaACCGGTATGAAGGGCAAGGAGGAGAAG |
|  | r1 | CACGCTGCAGcTGCTCCTCGATGTCC |
|  | f1 | GGACATCGAGGAGCAgCTGCAGCGTG |
|  | hCHIP reverse | GATATCaccctgaaaatacaaattctcGCTAGCTCAGTAGTCCTCCACCCAGC |
| FLAG-HSPA1A | HSPA1A forward | tccgaaaacctgtacttccaaggaaccggtatggccaaagccgcggcgatcggc |
|  | HSPA1A  reverse | atcaccctgaaaatacaaattctcgctagcctaatctacctcctcaatgg |
| HaloTag-ANXA5 | ANXA5 forward | CAGAGCGATAACGCGATCGCTTCCGAATTCatggcacaggttctcagagg |
|  | ANXA5 reverse | TCTAGATATCCGCGGTTGAGCTCTGAATTCttagtcatcttctccacag |
| **Target plasmid** | **Sequencing primer** | **Sequence** |
| EGFP-CHIP WT  EGFP-CHIP H260Q | FRTTO_For | tgacctccatagaagacacc |
|  | FRTTO_Rev | aactagaaggcacagtcgag |
|  | EGFP_F | catggtcctgctggagttcg |
|  | CHIP_For | atgaagggcaaggaggagaag |
| mCherry-CHIP WT  mCherry-CHIP H260Q  mCherry-CHIP K30A | mCherry_F | atcgtggaacagtacgaacg |
|  | FRTTO_Rev | aactagaaggcacagtcgag |
| FLAG-HSPA1A | FRTTO_For | tgacctccatagaagacacc |
|  | FRTTO_Rev | aactagaaggcacagtcgag |
|  | 19_106_s1 | tcaacgtgctgcggatcatc |
|  | 19_106_s2 | gagcatcaaccccgacgagg |
| HaloTag-ANXA5 | 24_212_s1 | aagaaaagtttatcaccatc |
|  | HaloTag_For | TCTGCTGCAAGAAGACAACC |
