## Supplementary material for "CHIP, VCP, and Nucleolar Gatekeepers Remodel the Nucleolus into a Stress-Responsive Proteostasis Hub": Table S5

| **SiRNA target** | **Source** | **Identifier** |
| --- | --- | --- |
| Silencer Select Negative control # 2 | Thermo Fisher Scientific | Cat# 4390846 |
| STUB1 | Thermo Fisher Scientific | Cat# 4392420 ID195025 |
| HSPA1A, HSPA1B | Thermo Fisher Scientific | Cat# AM16708 ID145248 |
| HSPA1A | Thermo Fisher Scientific | Cat# 4392420 IDs6965 |
| UTP15 | Thermo Fisher Scientific | Cat# 4392420 IDs38548 |
| NOLC1 | Thermo Fisher Scientific | Cat# 4392420 IDs17633 |
| NOL6 | Thermo Fisher Scientific | Cat# 4392420 IDs35200 |
| WDR18 | Thermo Fisher Scientific | Cat# 4392420 IDs33025 |
| WDR55 | Thermo Fisher Scientific | Cat# 4392420 IDs29591 |
| BRIX1 | Thermo Fisher Scientific | Cat# 4392420 IDs226839 |
| GNL3L | Thermo Fisher Scientific | Cat# 4392420 IDs29190 |
