## Supplementary material for "CHIP, VCP, and Nucleolar Gatekeepers Remodel the Nucleolus into a Stress-Responsive Proteostasis Hub": Table S6

| **Experiment** | **Compound** | **Compound treatment** | **Heat shock assay** |
| --- | --- | --- | --- |
| Nucleolar morphology and caps,  EGFP-CHIP migration | 0.05 µg/ml Actinomycin D | 30 min and 120 min pretreatment before HS | 90 min HS + 2 h recovery |
| Bystin immunofluorescence | 0.05 µg/ml Actinomycin D | Co-treatment with HS for 2 h | 2 h HS + 1-2 h recovery |
| HaloTag-RPL11 labeling in HeLa CHIP | 0.05 µg/ml Actinomycin D | 2 h pretreatment before HS | 90 min HS + 1 h recovery |
| FISH | 0.05 µg/ml Actinomycin D | 2 h pretreatment before HS | 90 min HS + 1 h recovery |
| FISH | 40 μM VER-155008 | 2 h pretreatment before HS | 90 min HS + 1 h recovery |
| Experiments in HEK293T | 40 μM VER-155008 | Co-treatment with HS for 2 h | 2 h HS + 1-2 h recovery |
| Experiment in MCF7 | 40 μM VER-155008 | Co-treatment with HS for 90 min | 90 min HS + 2 h recovery |
| Experiment in MCF7 | 40 μM VER-155008 | Co-treatment with Rec for 2 h | 90 min HS + 2 h recovery with the compound |
| EU labeling and EGFP-CHIP sequestration | 40 μM VER-155008 | 2 h pretreatment before HS | 90 min HS + 1 h HS with EU or 1 h recovery with EU |
| EU labeling and EGFP-CHIP sequestration | 100 μM SW02 | 3 h pretreatment before HS | 90 min HS + 1 h HS with EU or 1 h recovery with EU |
| EU labeling and EGFP-CHIP sequestration | 10 μM CB5083 | 2 h pretreatment before HS | 90 min HS + 1 h HS with EU or 1 h recovery with EU |
| EU labeling and EGFP-CHIP sequestration | 5 µg/ml Tunicamycin | 3 h pretreatment before HS | 90 min HS + 1 h HS with EU or 1 h recovery with EU |
| FBL and Ub staining in HeLa CHIP | 100 μM SW02 | 3 h pretreatment before HS | 90 min HS + 45 min recovery |
| FBL and Ub staining in HeLa CHIP | 40 μM VER-155008 | 2 h pretreatment before HS | 90 min HS + 45 min recovery |
| Ub, VCP and WDR55 IF in HeLa Flp | 10 μM CB5083 | 1 h pretreatment before HS | 90 min HS + 45 min recovery |
| Ub, CHIP IF upon siRNA treatment in HeLa Flp | 10 μM CB5083 | 1 h pretreatment before HS | 90 min HS + 45 min recovery |
| EGFP-CHIP sequestration, HaloTag-ANXA5 labeling, Ub, Bystin and FXR1 IF | 10 μM CB5083 | 2 h pretreatment before HS | 90 min HS |
| EGFP-CHIP sequestration and Bystin IF | 10 μM CB5083 | Co-treatment with Rec for 45 min | 90 min HS + 45 min recovery with the compound |
| Ub, VCP IF in HeLa Flp | 50 μM SMER28 | 1 h pretreatment before HS | 90 min HS + 45 min recovery |
| EGFP-CHIP sequestration, HaloTag-ANXA5 labeling, Ub, Bystin and FXR1 IF | 5 μM TAK243 | 2 h pretreatment before HS | 90 min HS |
| EGFP-CHIP sequestration, HaloTag-ANXA5 labeling, Ub and Bystin IF | 20 μM MG132 | Co-treatment with Rec for 45 min | 90 min HS + 45 min recovery with the compound |
| EGFP-CHIP sequestration | 5 μM MLN4924 | Co-treatment with Rec for 45 min | 90 min HS + 45 min recovery with the compound |
| EGFP-CHIP sequestration, HaloTag-ANXA5 labeling, Ub and Bystin IF | 50 μM PR619 | Co-treatment with Rec for 45 min | 90 min HS + 45 min recovery with the compound |
| EGFP-CHIP sequestration | 200 nM ISRIB | 30 min pretreatment before HS | 90 min HS + 45 min recovery |
| Bystin assay upon siRNA treatment | 10 nM Leptomycin B | Co-treatment with HS for 2 h | 2h HS |
| Bystin assay upon siRNA treatment | 10 nM Leptomycin B | Co-treatment with Rec for 1 h | 2h HS + 1 h recovery with the compound |
