## Supplementary material for "CHIP, VCP, and Nucleolar Gatekeepers Remodel the Nucleolus into a Stress-Responsive Proteostasis Hub": Table S7

| **Antibody target** | **Source** | **Identifier** | **Working dilution** |
| --- | --- | --- | --- |
| STUB1/CHIP | Abcam | Cat# ab134064 | 1:250 IF, 1:1000 WB |
| HSP70 | StressMarq Biosciences | Cat# SMC-100 | 1:250 IF, 1:500 WB |
| HSP70 | Abcam | Cat# ab181606 | 1:125 IF |
| NPM1 | Thermo Fisher Scientific | Cat# 32-5200 | 1:250 IF |
| NPM1 | Santa Cruz Biotechnology | Cat# sc-47725 | 1:200 IF |
| FBL | Cell Signaling Technology | Cat# 2639 | 1:400 IF, 1:500 WB |
| Alpha-tubulin | Thermo Fisher Scientific | Cat# 32-2500 | 1:1000 WB |
| Lamin B1 | Thermo Fisher Scientific | Cat# 33-2000 | 1:1000 WB |
| BYSL | Atlas Antibodies | Cat# HPA031219 | 1:300 IF |
| Mono- and polyubiquitinylated conjugates | Enzo Life Sciences | Cat# ENZ-ABS840 | 1:250 IF |
| VCP | Abcam | Cat# ab109240 | 1:250 IF |
| NOL6 | Atlas Antibodies | Cat# HPA055891 | 1:150 IF |
| NOL6 | Abcam | Cat# ab228836 | 1:120 IF |
| WDR55 | Novus Biologicals | Cat# NBP2-30625 | 1:200 IF |
| WDR55 | Atlas Antibodies | Cat# HPA048143 | 1:100 IF |
| FXR1 | Millipore | Cat# 03-176 | 1:250 IF |
| Alexa Fluor 488 goat anti-rabbit IgG | Molecular Probes |  | 1:500 |
| Alexa Fluor 568 goat anti-mouse IgG | Molecular Probes | Cat# A-11031 | 1:500 |
| Alexa Fluor 568 goat anti-rabbit IgG | Molecular Probes | Cat# A-11011 | 1:500 |
| Alexa Fluor 647 goat anti-mouse IgG | Molecular Probes | Cat# A-21235 | 1:500 |
| Alexa Fluor 647 goat anti-rabbit IgG | Molecular Probes | Cat# A-21245 | 1:500 |
| Cy3 AffiniPure Donkey Anti-Mouse IgG | Jackson ImmunoResearch | Cat# 715-165-150 | 1:1000 |
