## Supplementary material for "CHIP, VCP, and Nucleolar Gatekeepers Remodel the Nucleolus into a Stress-Responsive Proteostasis Hub": STAR Methods Table

| **REAGENT or RESOURCE** | **SOURCE** | **IDENTIFIER** |
| --- | --- | --- |
| **Antibodies (For detailed list of antibodies see also Table S7)** | | |
| Rabbit monoclonal anti-STUB1/CHIP | Abcam | Cat# ab134064, RRID:AB_2751008 |
| Mouse monoclonal anti-HSP70 | StressMarq Biosciences | Cat# SMC-100, RRID:AB_854199 |
| Rabbit monoclonal anti-HSP70 | Abcam | Cat# ab181606, RRID:AB_2910093 |
| Mouse monoclonal anti-NPM1 | Thermo Fisher Scientific | Cat# 32-5200, RRID:AB_2533084 |
| Mouse monoclonal anti-NPM1(FISH) | Santa Cruz Biotechnology | Cat# sc-47725, RRID:AB_628034 |
| Rabbit monoclonal anti-FBL | Cell Signaling Technology | Cat# 2639, RRID:AB_2278087 |
| Mouse monoclonal anti-Alpha-tubulin | Thermo Fisher Scientific | Cat# 32-2500, RRID:AB_2533071 |
| Mouse monoclonal anti-Lamin B1 | Thermo Fisher Scientific | Cat# 33-2000, RRID:AB_2533106 |
| Rabbit polyclonal anti-BYSL | Atlas Antibodies | Cat# HPA031219, RRID:AB_10601989 |
| - Mouse monoclonal anti-[Mono- and polyubiquitinylated conjugates](https://rrid.site/data/record/nif-0000-07730-1/AB_2935893/resolver?q=%2A&l=%2A&filter%5b%5d=Catalog%20Number:ABS840&i=rrid:ab_2935893-2871349) | Enzo Life Sciences | Cat# ENZ-ABS840, RRID:AB_2935893 |
| - Rabbit monoclonal anti-VCP | Abcam | Cat# ab109240, RRID:AB_10862588 |
| Rabbit polyclonal anti-NOL6 | Atlas Antibodies | Cat# HPA055891, RRID:AB_2682959 |
| Rabbit polyclonal anti-NOL6 | Abcam | Cat# ab228836, |
| Rabbit polyclonal anti-WDR55 | Novus Biologicals | Cat# NBP2-30625, RRID:AB_3277867 |
| Rabbit polyclonal anti-WDR55 | Atlas Antibodies | Cat# HPA048143, RRID:AB_2680282 |
| Mouse monoclonal anti-FXR1 | Millipore | Cat# 03-176, RRID:AB_10806485 |
| Alexa Fluor 488 goat anti-rabbit IgG | Molecular Probes | Cat# A-11034, RRID:AB_2576217 |
| Alexa Fluor 568 goat anti-mouse IgG | Molecular Probes | Cat# A-11031, RRID:AB_144696 |
| Alexa Fluor 568 goat anti-rabbit IgG | Molecular Probes | Cat# A-11011, RRID:AB_143157 |
| Alexa Fluor 647 goat anti-mouse IgG | Molecular Probes | Cat# A-21235, RRID:AB_2535804 |
| Alexa Fluor 647 goat anti-rabbit IgG | Molecular Probes | Cat# A-21245, RRID:AB_2535813 |
| Cy3 AffiniPure Donkey Anti-Mouse IgG | Jackson ImmunoResearch | Cat# 715-165-150, RRID:AB_2340813 |
| **Chemicals, peptides, and recombinant proteins** | | |
| Dimethyl sulfoxide | Sigma-Aldrich | Cat# D4540 |
| MLN4924 | Selleckchem | Cat# S7109 |
| Leptomycin B | Sigma-Aldrich | Cat# L2913 |
| Actinomycin D | Tocris | Cat# 1229 |
| MG132 | Selleckchem | Cat# S2619 |
| PR-619 | Life Sensors | Cat# SI9619 |
| TAK-243 | Selleckchem | Cat# S8341 |
| CB-5083 | MedChemExpress | Cat# HY-12861 |
| SMER28 | Tocris | Cat# 4297 |
| Tetracycline HCL | BioShop | Cat# TET701 |
| SW02 | Sigma-Aldrich | Cat# S5951 |
| VER-155008 | Sigma-Aldrich | Cat# SML0271 |
| Tunicamycin | Santa Cruz Biotechnology | Cat# sc-3506 |
| 7-Bromo-1-heptanol | VWR | Cat# H54762.03 |
| TRI Reagent | Sigma-Aldrich | Cat# 93289 |
| No-Stain Protein Labeling Reagent | Thermo Fisher Scientific | Cat# A44449 |
| Lipofectamine 2000 Transfection Reagent | Thermo Fisher Scientific | Cat# E2312 |
| Lipofectamine RNAiMAX Reagent | Thermo Fisher Scientific | Cat# 13778150 |
| TransIT-HeLaMONSTER Transfection Kit | Mirus | Cat# MIR 2900 |
| TransIT-293 Transfection Kit | Mirus | Cat# MIR 2700 |
| Janelia Fluor 646 HaloTag Ligand | Promega | Cat# GA1121 |
| Hoechst 33342 | Thermo Fisher Scientific | Cat# 62249 |
| DAPI | Sigma-Aldrich | Cat# D9542 |
| Paraformaldehyde solution 4% in PBS (FISH) | Santa Cruz Biotechnology | Cat# sc-281692 |
| PerfectHyb Plus Hybridization Buffer | Sigma-Aldrich | Cat# H7033 |
| 20x SSC Buffer | Thermo Fisher Scientific | Cat# 15557044 |
| 16% Formaldehyde methanol-free | Thermo Fisher Scientific | Cat# 28906 |
| Hygromycin B | Thermo Fisher Scientific | Cat# 10687010 |
| Blasticidin S | Invivogen | Cat# ant-bl-05 |
| G418 | Thermo Fisher Scientific | Cat# 10131035 |
| Vectashield Vibrance Antifade Mounting medium with DAPI | Vector Laboratories | Cat# H-1800-10 |
| Protease inhibitor cocktail | Roche | Cat# 04 693 159 001 |
| Me4BodipyFL-Ahx3Leu3VS Proteasome Activity Probe | Bio-Techne | Cat# I-190 |
| Poly-L-lysine | Sigma-Aldrich | Cat# P4707 |
| **Critical commercial assays** | | |
| Click-iT™ Plus OPP Alexa Fluor™ 488 Protein Synthesis Assay Kit | Thermo Fisher Scientific | Cat# 10456 |
| Click-iT™ Plus OPP Alexa Fluor™ 594 Protein Synthesis Assay Kit | Thermo Fisher Scientific | Cat# 10457 |
| Click-iT™ RNA Alexa Fluor™ 594 Imaging Kit | Thermo Fisher Scientific | Cat# 10330 |
| SuperSignal West Pico PLUS Chemiluminescent Substrate | Thermo Fisher Scientific | Cat# 34580 |
| Pierce Rapid Gold BCA Protein Assay Kit | Thermo Fisher Scientific | Cat# A53225 |
| **Deposited data** | | |
| RNA sequencing raw data | This study | GSE190142 |
| **Experimental models: Cell lines** | | |
| Human: HeLa Flp-In T-REx | Szczesny et al. | N/A |
| Human: HEK293 Flp-In T-REx | Szczesny et al. | N/A |
| Human: MCF7 | Zylicz et al. | N/A |
| Human: HEK293T NLS LG | Hipp et al. | N/A |
| Human: HeLa CHIP | This study | N/A |
| Human: HEK CHIP | This study | N/A |
| Human: HeLa EGFP-NPM1 | Szczesny et al | N/A |
| **Oligonucleotides** | | |
| See Table S5 for siRNA list | N/A | N/A |
| See Table S4 for primer list | N/A | N/A |
| rRNA FISH probe  47S-rRNA: 5'-CGGAGGCCCAACCTCTCCAGCGACAGGTCGCCAGAGGACAGCGTGTCAGC-Cy5 | Szaflarski et al.  Custom synthesis performed at the Institute of Biochemistry and Biophysics, PAS, Warsaw | N/A |
| **Recombinant DNA** | | |
| pKK-EGFP-TEV | Szczesny et al. | N/A |
| pKK-mCherry-TEV | Szczesny et al. | N/A |
| pKK-FLAG-TEV | Szczesny et al. | N/A |
| pKK-EGFP-CHIP | This study | N/A |
| pKK-EGFP-CHIP H260Q | This study | N/A |
| pKK-mCherry-CHIP | This study | N/A |
| pKK-mCherry-CHIP H260Q | This study | N/A |
| pKK-mCherry-CHIP K30A | This study | N/A |
| pKK-FLAG-HSPA1A | This study | N/A |
| pOG44 | Szczesny et al. | N/A |
| pHTN-HaloTag CMV-neo | Promega | Cat# G7721 |
| pFN21A-HaloTag-RPL11 | Promega | Cat# FHC02105 |
| pHTN-HaloTag-ANXA5 | This study | N/A |
| **Software and algorithms** | | |
| GraphPad Prism 9 (v. 9.5.0) | Prism | <https://www.graphpad.com>, RRID:SCR_002798 |
| ImageJ | Schindelin et al. | <https://imagej.net/software/fiji/>, RRID:SCR_003070 |
| Inkscape | N/A | RRID:SCR_014479 |
| BioRender | N/A | <https://www.biorender.com/>, RRID:SCR_018361 |
| CellProfiler 4.2.1 | Stirling et al. | <https://cellprofiler.org/>, RRID:SCR_007358 |
| FRAPAnalyser | N/A | https://github.com/ssgpers/FRAPAnalyser |
| Magellan | Tecan | https://lifesciences.tecan.com/software-magellan |
| R | N/A | https://www.r-project.org/ |
| Imaris 7.4.2 | Oxford Instruments | <https://imaris.oxinst.com/>, RRID:SCR_007370 |
| **Other** | | |
| Dulbecco′s Modified Eagle′s Medium | Sigma-Aldrich | Cat# D6429 |
| Fetal Bovine Serum | Sigma-Aldrich | Cat# F9665 |
| Fetal Bovine Serum Tet-system approved | Thermo Fisher Scientific | Cat# A4736201 |
| Antibiotic-Antimycotic | Thermo Fisher Scientific | Cat# 15240062 |
| Opti-MEM I Reduced Serum Medium | Thermo Fisher Scientific | Cat# 11058021 |
| Goat serum | Sigma-Aldrich | Cat# G9023 |
| Trypsin-EDTA solution | Sigma-Aldrich | Cat# T4049 |
| DPBS | Sigma-Aldrich | Cat# D8662 |
| µ-Slide 8 Well | Ibidi | Cat# 80826 |
| Glass Bottom Dish 35 mm | Ibidi | Cat#8156 |
